# Host shifting and host sharing in a genus of specialist flies diversifying alongside their sunflower hosts

**DOI:** 10.1101/2020.03.18.995589

**Authors:** Alaine C. Hippee, Marc A. Beer, Robin K. Bagley, Marty A. Condon, Andrew Kitchen, Edward A. Lisowski, Allen L. Norrbom, Andrew A. Forbes

**Affiliations:** Department of Biology, University of Iowa, Iowa City, IA 52242, USA; School of Biological Sciences, Washington State University, Pullman, WA 99164, USA; Department of Biology, Cornell College, Mount Vernon, IA 52314, USA; Department of Anthropology, University of Iowa, Iowa City, IA 52242, USA; Washington Department of Agriculture, Yakima, Washington, 98902 USA; Systematic Entomology Laboratory, USDA, ARS, PSI, c/o National Museum of Natural History, Washington, DC 20013 USA

## Abstract

Congeneric parasites are unlikely to specialize on the same tissues of the same host species, likely because of strong multifarious selection against niche overlap. Exceptions where multiple congeneric species overlap on the same tissues may therefore reveal important insights into the ecological factors underlying the origins and maintenance of diversity. Larvae of sunflower maggot flies in genus *Strauzia* feed on the pith of plants in the family Asteraceae. Although *Strauzia* tend to be host specialists, some species overlap in their host use. To resolve the origins of host sharing among these specialist flies, we used reduced representation genomic sequencing to infer the first multi-locus phylogeny of genus *Strauzia.* Our results show that *Helianthus tuberosus* and *Helianthus grosseserratus* each host three different fly species, and that the flies co-occurring on a host are not one another’s closest relatives. Though this pattern implies that host sharing is most likely the result of host shifts, these may not be host shifts in the conventional sense of an insect moving onto an entirely new plant. Many hosts of *Strauzia* belong to a young (1-2 MYA) clade of perennial sunflowers noted for their frequent introgression and hybrid speciation events. In at least one case, flies may have converged upon a host after their respective ancestral host plants hybridized to form a new sunflower species (*H. tuberosus*). Broadly, we suggest that rapid and recent adaptive introgression and speciation in this group of plants may have instigated the diversification of their phytophagous fly associates, including the convergence of >1 species onto the same shared host plants.

## Introduction

Selection is expected to disfavor use of the same host niche by multiple, closely related specialist parasite species, primarily due to competitive exclusion and other forms of interspecific conflict (Miller 1967; Denno et al. 1995; Berlocher 1998; Craig et al. 2000). This principle generally holds true for parasitic insects, the most species-rich of all animals (Janz et al. 2006; Futuyma and Agrawal 2009): competition for resources (Jaenike 1990; Denno et al. 1995; Abrahamson et al. 1993; Violle et al. 2011), reproductive conflict (Funk 1998; Nosil et al. 2003; Groning and Hochkirch 2008), and avoidance of habitat-specific predators (Bernays and Graham 1988; Ohsaki and Sato 1994) should all favor the evolution of reduced overlap in host use.

Host use is also relevant to the origins of new specialist insect species. When insects shift onto new hosts, accompanying changes in behavior and life history can result in the evolution of reproductive isolation between different host-associated populations (Bush 1969; Funk 1998; Via 1999; Dres and Mallet 2002; Matsubayashi et al. 2010). In some cases, host shifts have been directly implicated in the origin of new incipient species (Feder et al. 1988; Sheldon and Jones 2001; Carroll et al. 2005; Matsubayashi et al. 2013; Hood et al. 2015; Forbes et al. 2017). Host shifting often represents a move into a new and relatively unoccupied habitat, such that competition for space and resources is reduced (Feder et al. 1995; Nosil 2012). This advantage is important because host shifts are associated with many other physiological, ecological, and evolutionary constraints (Hardin 1960; Menken 1995; Silva-Brandao and Solferini 2007).

Considering that previous theoretical and empirical work addressing host sharing and host shifting agrees that congeneric insect species should not often co-occur, it is therefore surprising that in some genera multiple species specialize on the same tissue of the same plant species in sympatry. In the Neotropical tephritid genus *Blepharoneura*, the larvae of several closely related species are often found feeding on the same calyx tissues of the same cucurbit vines (Condon et al. 2008, 2014; Winkler et al. 2018). Similarly, cecidomyiid gall flies have undergone several within-host adaptive radiations, resulting in multiple morphs utilizing the same host plants (Joy and Crespi 2007; Stireman et al. 2008). Understanding how such situations arise – and how they persist – can reveal important new insights into the factors governing the origin and maintenance of specialist insect diversity.

Sunflower maggot flies in the genus *Strauzia* (Diptera: Tephritidae) represent one of these unusual cases where multiple specialist congeneric fly species may have shifted onto the same host plant tissue. *Strauzia* are associated with North American plants in the family Asteraceae, and their life cycles, as in most tephritids, are intimately linked to their plant hosts. Flies meet and mate on their host plant’s leaves, and females oviposit into the upper nodes of the plant stem (Stoltzfus 1988). Upon hatching, larvae feed on the plant’s pith, mining down toward the roots where they eventually pupate either in the root crown or the soil around the plant. Most *Strauzia* species associate with a single host species, and, in turn, most hosts harbor only one fly species (Steyskal 1986; Stoltzfus 1988). However, decades of accumulated evidence suggest that the sawtooth sunflower (*Helianthus grosseserratus*) and Jerusalem artichoke (*H. tuberosus*) are each host to multiple *Strauzia* species. Either two or three fly species are associated with *H. grosseserratus* (Lisowski 1985; Steyskal 1986; Stoltzfus 1988), while *H. tuberosus* hosts three genetically differentiated lineages that have variably been referred to as varieties or species, depending on the author (Stoltzfus 1988; Forbes et al. 2013; Hippee et al. 2016). Evaluating how multiple *Strauzia* species have come to coexist on their respective plants promises exciting new insights into the origins and maintenance of insect diversity. However, such work currently has two major impediments: *Strauzia* taxonomy remains unresolved, and – critical to unraveling histories of host shifting – the phylogeny of the genus is unknown.

*Strauzia* taxonomy has a complex history, with both the number and identity of species being subjects of disagreement. All species were initially lumped under one name, *Trypeta longipennis* Wiedemann, 1830 (currently *Strauzia longipennis* (Wiedemann)), but additional divisions based on both fly morphology and host plant affiliation quickly became evident. In his treatment of North American Diptera, Loew (1873) states: “It cannot be doubted that [*Strauzia*] *longipennis* Wied. either is a very variable species, or that North America possesses a number of closely allied species…” Loew himself recognized seven “varieties” of *S. longipennis*, but recognized none of these as species.

Taxonomic work in the latter half of the 20th century split Loew’s single species into many, but different authors came to different conclusions as to how many species warranted recognition, as well as the identities of those species (Table 1). First, Lisowski (1985) recognized six of Loew’s varieties as species, and added one additional species using a manuscript name (“*S. bushi*”) that has never been published and was largely overlooked by subsequent authors; we therefore refer to it as “Bush’s Fly” (note that we use quotation marks when we refer to this fly because “Bush’s Fly” is not an available name). The following year, Steyskal (1986) also recognized three of Loew’s varieties as species, adding two more species, *Strauzia verbesinae* and *Strauzia stoltzfusi*. Two years later, Stoltzfus (1988) revised the genus again, recognizing all but one of Loew’s remaining varieties as species (the exception was *S. longipennis* var. *confluens*, which Loew (1873) had described based on a single male). Stoltzfus (1988) also added three new species: *S. rugosum*, *S. uvedaliae*, and *S. noctipennis*. An appropriate summary of the recent state of *Strauzia* taxonomy comes from Foote et al. (1993), who, regarding many of the species named above, say: “…we have no proof at this time that those taxa, based as they are mainly on host preference, are deserving of full species status.”

**Table 1.**
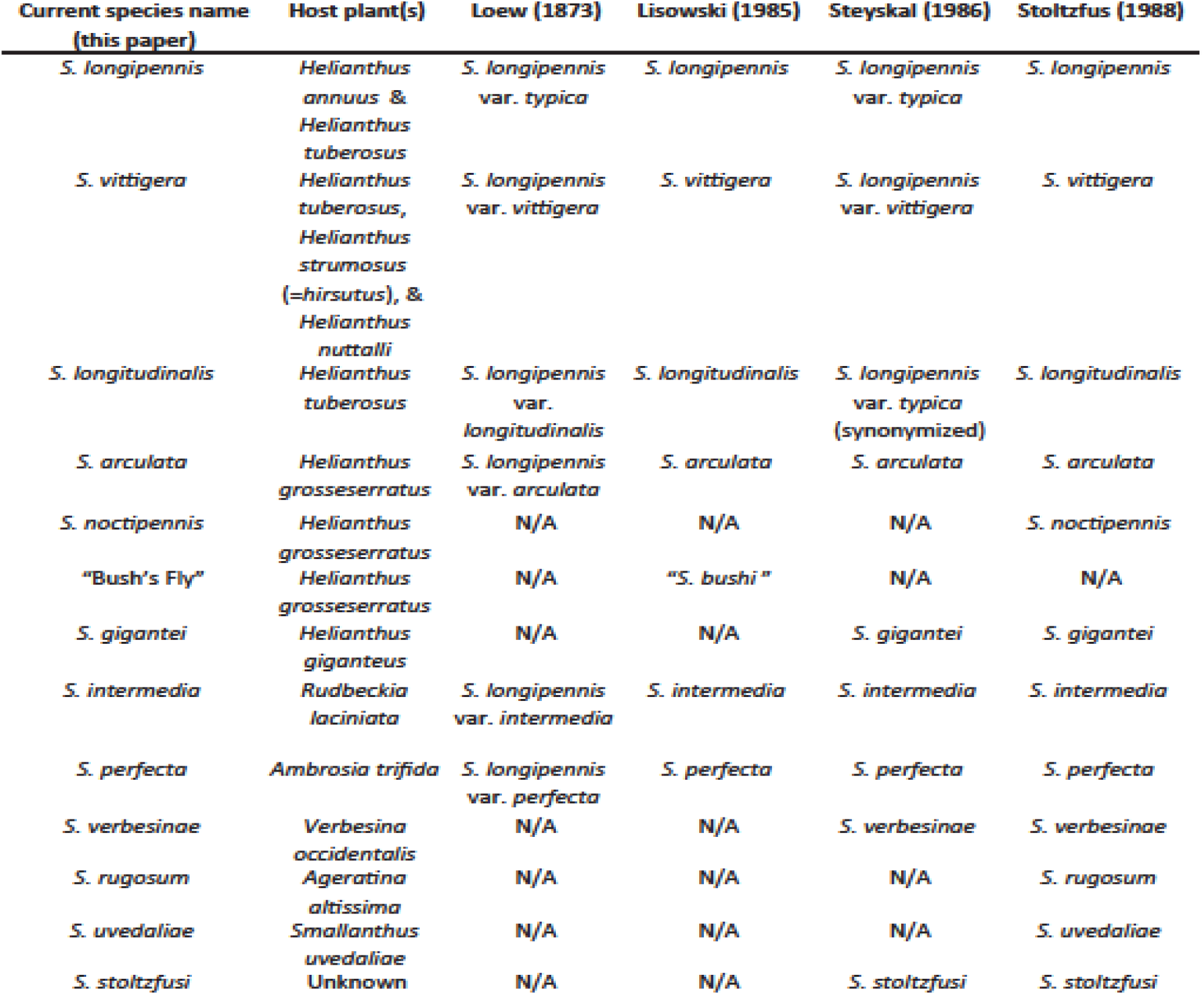
Summary of historical *Strauzia* species names. Loew’s (1873) variety *confluens* is omitted from this table, as it was described from a single male fly and subsequent authors have tended to ignore it. The enigmatic *S. stoltzfusi* was not collected for this study and its status is unknown. N/A refers to species that were not mentioned in the paper.

Disagreement among authors has perhaps been most pronounced in dealing with putatively distinct *Strauzia* species that may share the same plant hosts. Alongside *S. arculata*, Stoltzfus’ (1988) *S. noctipennis* and Lisowski’s (1985) “Bush’s Fly” are all described as flies associated with *H. grosseserratus*, but it is not clear whether “Bush’s Fly” and *S. noctipennis* are synonyms or different species. Even more troublesome is the complex of *Strauzia* flies associated with *H. annuus*, *H. tuberosus*, and other related sunflowers (Table 1). Most authors agree that they may represent several different taxa, but the range of morphological variation is such that characters have not yet emerged that definitively delimit any one species from another, and they have oscillated between being called “species” and “varieties.”

Genetic approaches have helped address this confusion. Axen et al. (2010) made some initial progress, identifying two haplotype groups within the *S. longipennis* complex, each closely associated with a particular thoracic striping pattern – though the association between haplotype and morphology was imperfect. Later, Forbes et al. (2013) and Hippee et al. (2016) used AFLP and microsatellite markers, respectively, to demonstrate that the *Strauzia* flies reared from *Helianthus tuberosus* include three genetically distinct groupings. The microsatellite-based analyses in particular showed a 1:1 correlation between each genetic cluster and a particular combination of thoracic stripes and wing patterns (Hippee et al. 2016). This work strongly suggests that the *H. tuberosus*-associated *S. longipennis* complex consists of at least three different species, but their relationships to one another, and to flies with similar morphologies on other host plants, remains uncertain.

The apparent coexistence of more than one specialist *Strauzia* species on each of *H. grosseserratus* and *H. tuberosus* is sufficiently novel among specialist parasitic insects that we seek to understand their respective origins. To this end, two primary hypotheses could explain how *Strauzia* have come to coexist on the same plant. First, fly species sharing a single host may all share a recent common ancestor that used the same plant host, and which diversified without host shifts, perhaps during periods of geographic isolation (e.g., in refuges during glacial periods; Rand 1948). In this “no host shift” scenario (proposed in Hippee et al. 2016), the now-reproductively isolated sister species would later have become sympatric once the temporarily fragmented geographic distribution of their host plant fused back into a contiguous range. An alternative “host shift” scenario is that flies originating on different hosts converged upon single plant species via multiple host shifting events. These alternative scenarios can be resolved with a phylogeny. A “no host shift” hypothesis would be supported by a tree that shows common ancestry among flies using the same host plant, while for a “host shift” hypothesis, flies currently using the same host plant should be more closely related to species using other host plants than they are to one another. These hypotheses are also not mutually exclusive; two species may diverge from one another without changing hosts, while a third joins them after a host-shift.

In this study, we use wide-ranging collections of *Strauzia* from many different plant hosts and geographic locations, a reduced representation genomic sequencing approach, and phylogenetic inference to evaluate the following hypotheses: 1) *H. tuberosus* and *H. grosseserratus* are each host to three genetically distinct *Strauzia* species; and 2) *Strauzia* species sharing the same host plant did not diverge while on their respective host plants, but instead are the result of independent host shifts from other plant hosts. We also use the *Strauzia* phylogeny to consider how known patterns of evolution and speciation of North American sunflowers may have affected recent trajectories of *Strauzia* speciation.

## Methods

### Sample collection and DNA extraction

We collected *Strauzia* flies as adults, larvae, and pupae from a diverse set of geographic locations and plant hosts across North America (Figure 1; Figure S1; Table S1). Collections represented all named *Strauzia* species except for *S. stoltzfusi* (Steyskal 1986). We captured adults individually in plastic cups on their plant hosts, preserved them in 95% ethanol, and identified them to putative species using morphological diagnoses suggested by Lisowski (1985), Steyskal (1986), and Stoltzfus (1998). Larvae and pupae, which are not identifiable to species based on morphological characteristics, were collected by uprooting known host plants, carefully bisecting plant stems, and removing the immature flies within. First- and second-instar larvae were immediately preserved in 95% ethanol. A subset of third instar larvae were maintained until pupation in moist vermiculite and then overwintered in a refrigerator for 4 months at 4-8°C. We also overwintered all *Strauzia* pupae collected from host plants in these same conditions. After 4 months, we removed pupae from the refrigerator, held them at 18°C for 1 week, and then moved them to a light- and temperature-controlled incubator (16:8 photoperiod; 25°C) to encourage post-diapause development and eclosion of adults. All ethanol-preserved *Strauzia* were stored at −80°C prior to DNA extraction.

**Figure 1.**
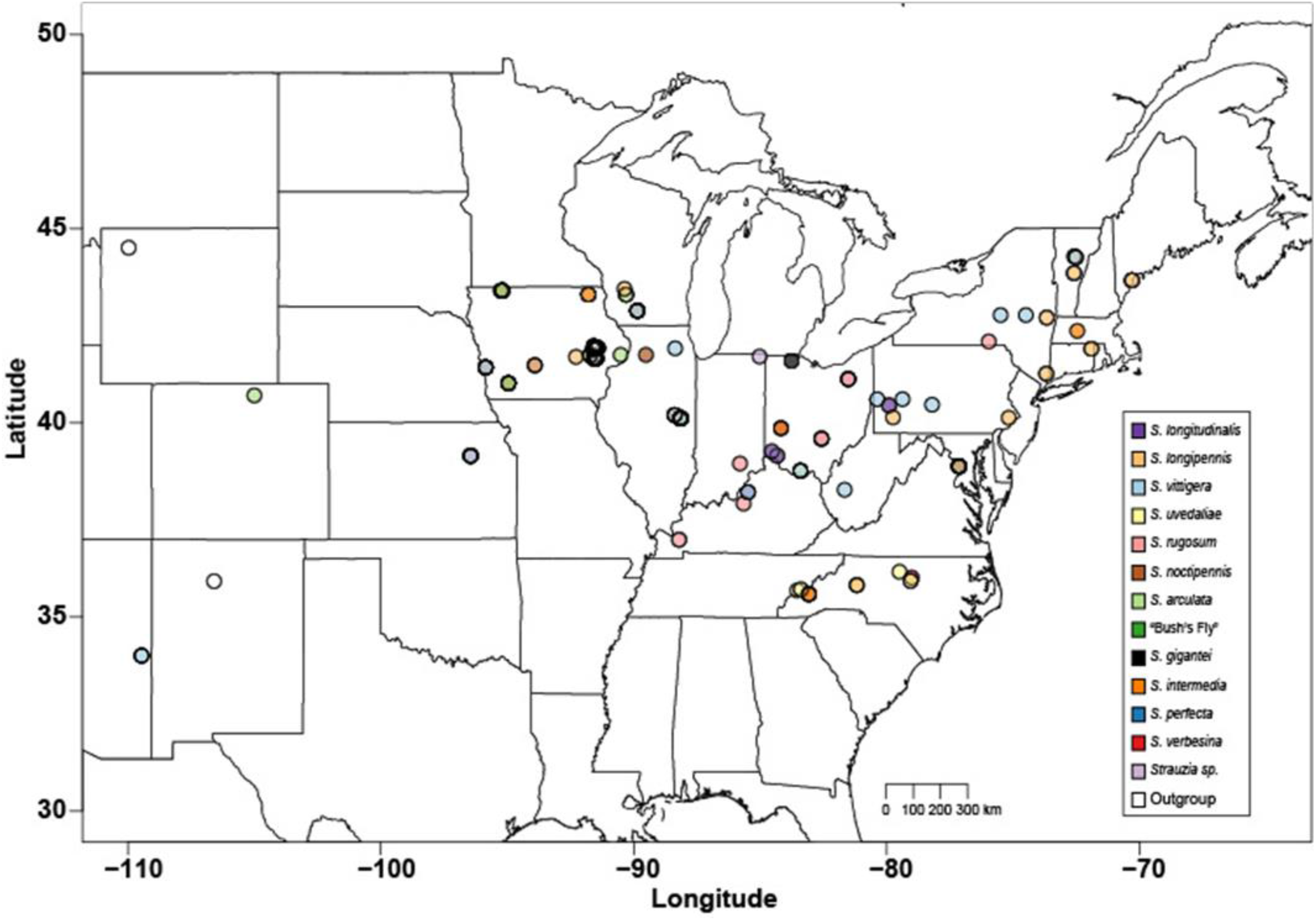
Map showing the collection location of all *Strauzia* samples used in the mtDNA and SNP phylogenetic analyses. Colored points correspond to species identifications described in the legend. See Table S1 for collection information, including host plant identities, GPS locations, whether each sample was used in the mtCOI or 3RAD datasets, or in both.

We extracted DNA from 205 preserved *Strauzia* flies using either a Qiagen Blood and Tissue Kit (Qiagen Sciences, Germantown, Maryland) or a modified CTAB extraction protocol based on Chen et al. (2010). We also extracted DNA from various other tephritid species (*Chetostoma rubidium*, *Paramyiolia rhino*, *Rhagoletis pomonella*, *Trypeta flaveola*, *Trypeta tortilis*, and *Zonosemata electa*) for use as outgroups. Each sample was then visualized on a 0.8% agarose gel to assess the degree of DNA degradation. The subset of individuals used in the 3RAD library prep (see below) were also quantified using the fluorescence-based Quant-iT PicoGreen dsDNA Assay Kit (Invitrogen - Molecular Probes, Eugene, OR, USA). Extracted DNA were stored at −20°C until use.

### MtCOI sequencing and phylogenetic analysis

We constructed a mitochondrial cytochrome oxidase I (COI) gene tree as a preliminary assessment of genetic relationships among species. For a subset of 88 *Strauzia* and 5 individuals from three outgroup species (*T. flaveola*, *C. rubidium*, and *P. rhino*), we amplified 627 bp of the “barcoding” region of the COI gene using the primers LepF1 and LepR1 (Smith et al. 2008). We PCR amplified COI using the following thermocycler program: 94°C for 3 min, followed by 35 cycles of 94°C for 30 seconds, 46°C for 1 minute, and 72°C for 2 minutes, and a final extension step of 72°C for 4 minutes, as in Forbes et al. (2013). All samples were checked for successful COI amplification using gel electrophoresis. We cleaned samples in preparation for sequencing using Shrimp Alkaline Phosphatase (USB, Swampscott, MA) and Exonuclease I (New England Biolabs, Ipswich, Maryland) and cycle sequenced in the forward and reverse direction using an ABI 3730 DNA Sequencer (Applied Biosystems, Waltham, Massachusetts) and the Big Dye Terminator v3.1 Cycle Sequencing Kit (Applied Biosystems, Waltham, Massachusetts). Forward and reverse reads were inspected visually and assembled into consensus sequences using SeqMan (DNASTAR, Madison, WI). We aligned the consensus sequences of all 93 individuals using ClustalW (v1; Thompson et al. 1994) implemented in BioEdit (v7.2.5; Hall 1999) and model-tested using jModelTest2 (v2.1.7; Darriba et al. 2012; Guindon and Gascuel 2003). The Bayesian phylogenetic analysis was computed using MrBayes (v.3.1.2; Huelsenbeck and Ronquist 2003) with a GTR + Γ substitution model, as recommended by the AIC criterion in jModelTest2 and 1000 bootstrap replicates. We visualized the phylogeny using FigTree (v1.4.3; Rambaut 2007) (Figure S2).

### 3RAD library preparation and sequencing

We also generated a multilocus single nucleotide polymorphism (SNP) dataset using a restriction site associated DNA (RAD) sequencing approach. In sets of up to 40 individuals, sorted by yield and randomized by location and host plant, we prepared RAD libraries for a total of 164 individuals (162 *Strauzia* plus two flies from an outgroup species, *Zonosemata electa*) following the “3RAD” protocol outlined in Bayona-Vásquez et al. (2019). Briefly, genomic DNA was digested with the restriction enzymes NheI, XbaI, and EcoRI for 1 hour at 37°C. Immediately following digestion, we ligated a unique combination of barcoded adapters to each sample (Table S2) and removed adapter dimers by alternating between conditions favoring ligation (20 minutes at 22°C) and digestion (10 minutes at 37°C) twice. The restriction enzymes were then heat-killed. We pooled the barcoded samples and labeled each pool with one of four iTru7 Illumina multiplexing indexes and a universal iTru5 Illumina index (Table S2; Bayona-Vásquez et al. 2019) via 15 rounds of high-fidelity PCR amplification (KAPA HotStart PCR Kit, KAPA Biosystems, Wilmington, Massachusetts). Each pooled, amplified library was visualized on an agarose gel to confirm successful library creation and then loaded onto a BluePippin (Sage Science, Beverly, MA) for automated size-selection of 550±50bp fragments. Final libraries were fluorescently quantitated (Quant-iT PicoGreen dsDNA Assay Kit, Invitrogen, Carlsbad, CA), pooled, and sequenced using 150bp paired-end reads on an Illumina HiSeq 4000 sequencer (Illumina, San Diego, CA) at the Iowa Institute of Human Genomics at the University of Iowa.

### Demultiplexing and SNP calling

Raw reads were demultiplexed based on their iTru7 index by the bioinformatics division at the Iowa Institute of Human Genomics. We further demultiplexed the reads based on their inline barcode combination using the *process_RADtags* pipeline for paired-end reads implemented in Stacks2 (v2.4; Rochette et al. 2019). We also used this pipeline to perform initial quality control, including removal of sequences with a Phred score lower than 20, and trimmed any reads contaminated by the universal Illumina adapter sequence. Following demultiplexing, we discarded all R2s as they are physically linked to the R1s.

We next generated SNP datasets by processing the filtered, demultiplexed reads through iPyrad (v0.7.30; Eaton and Overcast 2016) with a maximum clustering threshold of 0.85 and a minimum depth for base calls for both statistical and majority rule set to 10. A preliminary maximum likelihood phylogenetic analysis using RAxML (v8.2.12; Stamatakis 2014) including just the outgroup and a subset of 13 *Strauzia* individuals representing seven species supported the general basal topology resolved from mtCOI data (Figure S3), and importantly offered a second line of support for *S. perfecta* being the most basal *Strauzia* species. We then generated a final SNP dataset that included all 127 *Strauzia* individuals that passed the initial filtering steps and excluded the two *Z. electa* individuals in order to maximize the number of SNPs and loci available for the phylogenetic analysis within *Strauzia*. The final dataset includes a maximum of 20 SNPs per locus and up to 70% missing data per site. All of the subsequent phylogenies were rooted using *S. perfecta*, an approach justified by the agreement between the mtCOI phylogeny and the preliminary 3RAD analysis.

### Phylogenetic analysis of SNP datasets

Using the complete *Strauzia* SNP dataset, we generated phylogenies using maximum likelihood, Bayesian, and quartet-based approaches to account for methodological variation in outcomes when processing SNP data (Ree and Hipp 2015; Leache and Oakes 2017). We generated a maximum likelihood phylogeny using RAxML (v8.2.12; Stamatakis 2014) with a GTR+Γ substitution model and 1000 bootstrap replicates. The Bayesian analysis tree was generated in MrBayes (v3.2.6; Ronquist and Huelsenbeck 2003) using a GTR+Γ substitution model and 1000 bootstrap replicates. We used SVDQuartets (Chifman and Kubatko 2014; Chifman and Kubatko 2015) implemented in PAUP* (v4.0a166; Swofford 2001) to infer a quartet-based coalescent tree, which allows for missing data in SNP matrices and assumes each SNP has an independent evolutionary history (Leache and Oaks 2017). We calculated all possible quartets from our *Strauzia* SNP matrix and performed 1000 bootstrap replicates. The quartet-based phylogeny was visualized using FigTree (v1.4.3; Rambaut 2007), and the maximum likelihood and Bayesian phylogenies were visualized together using TreeGraph2 (v2.15.0; Stover and Muller 2010), with the Bayesian posterior probabilities overlaid on the maximum likelihood phylogeny on all nodes that were supported using both phylogenetic methods.

### Phylogenetic discordance and introgression

We looked for evidence of discordance between gene trees and the final species tree, as concatenated datasets can converge on an incorrect topology as a result of statistical inconsistency in the coalescent model (Wagner et al. 2013; Ree and Hipp 2015). By generating phylogenetic trees from each locus individually, we can quantify the amount of phylogenetic discordance among loci to determine the phylogenetic support for a population tree using the program BUCKy (v1.4.4; Ane et al. 2007; Larget et al. 2010), which assumes incomplete lineage sorting is included in the dataset. For this analysis, we generated a full-locus dataset in iPyrad (Eaton and Overcast 2016) that included one individual from each *Strauzia* clade to reduce computation time. We focused our analysis on the *Strauzia* species that were not resolved in the mtCOI phylogenetic analysis, excluding both *S. perfecta* and *S. intermedia*, as well as all clades that were represented by a single individual. BUCKy does not permit missing data, so we included only those loci that were shared across all 8 included *Strauzia* samples. Using this whole-locus dataset, we generated a phylogeny for each locus individually using MrBayes (v3.2.6; Ronquist and Huelsenbeck 2003) and a GTR+Γ substitution model for 1,000,000 generations. We used *mbsum*, implemented in BUCKy, to summarize the posterior distributions of each locus analysis and then calculated concordance factors with four chains for 500,000 generations. We repeated the analysis with α, the *a priori* gene discordance parameter, set at 1, 10, and 100. Both the population tree and primary concordance tree were visualized using FigTree (v1.4.3; Rambaut 2007).

Because some *Strauzia* species co-occur on the same plant hosts, we looked for evidence of introgression by implementing Patterson’s D-statistic tests (Durand et al. 2011) in iPyrad (Eaton and Overcast 2016). We generated a subset of species comparisons using our maximum likelihood phylogenetic tree with the constraint that *S. perfecta* was the outgroup in all analyses. Introgression analyses were not possible between all species pairs, as some phylogenetic relationships violate the assumptions of the Patterson’s D-statistic test, but we tested for introgression among some fly species found on the same plant hosts, as well as between close relatives using different hosts. Each analysis was repeated for all possible combinations with 1000 bootstrap replicates. The D-statistics and associated Z-scores were summarized across each clade and test in the analysis.

## Results

### Phylogenetic inference of COI

The mtCOI phylogeny of *Strauzia* included 627 base pairs from 93 individuals (Figure S2). Flies putatively identified by morphology and/or host plant use (when collected as larvae) as being *S. arculata*, *S. intermedia*, *S. verbesinae*, and *S. perfecta* formed distinct clades, but all other species (including *S. noctipennis*, *S. longipennis*, *S. vittigera*, *S. longitudinalis*, *S. gigantei*, *S. rugosum*, “Bush’s Fly”, and *S. uvedaliae*) formed a large polytomy. Bootstrap support was high among the most basal clades, including *S. perfecta*, *S. verbesinae*, *S. intermedia*, and *S. arculata*. However, support throughout the rest of the tree was variable, with relatively low support throughout its internal nodes.

### Phylogenetic inference from SNP datasets

The 3RAD sequencing generated an average of 2.8 million reads per individual. After quality filtering in Stacks2 and iPyrad, we retained an average of 1.6 million (±1.3 million SD) reads per individual. We excluded 35 individuals that retained less than 47,500 reads, as well as the two *Z. electa* individuals. The resulting dataset included an average of 539 loci per individual, with a total of 1325 unique loci and 17514 SNPs. This dataset allowed up to 70% missing data at any given locus, with an average of 58.8% missing data for individuals included in the phylogenetic analysis.

All three phylogenies (Maximum Likelihood, Bayesian, and quartet-based coalescent methods) supported monophyly for each of the 12 known *Strauzia* species included in this study. Maximum likelihood and Bayesian phylogenies (Figure 2) show support for the same topology among the species of *Strauzia*, with *S. perfecta*, *S. verbesinae*, and *S. intermedia* as the most basal clades. Statistical support was high in most nodes across both trees with the majority of internodes having bootstrap support >98 and/or posterior probabilities >0.95. In the Bayesian phylogeny, the node separating *S. verbesinae* and *S. intermedia* had a lower posterior probability (0.73), possibly as a result of the *S. verbesinae* clade being represented by a single individual. Statistical support for both trees was notably lower at the tips, with many nodes within *Strauzia* species clades falling below the 50% threshold. The quartet-based phylogeny (Figure S4) differs from the Maximum Likelihood and Bayesian phylogenies in the placement of *S. arculata* and *S. verbesinae*, but otherwise retains the same relationships among the *H. tuberosus*-associated *Strauzia* species. These topological differences may be a result of bioinformatic processing, differences among phylogenetic programs in the handling of missing data, or incomplete lineage sorting (Leache et al. 2015). The placement of *S. arculata* in the phylogenetic trees generated using Maximum Likelihood and Bayesian approaches (Figure 2) is supported by the gene discordance analysis (discussed below), but also suggests that there are many conflicting topologies regarding the placement *S. arculata*. Only one *S. verbesinae* individual was included in these phylogenetic analyses, so we cannot determine which topology is more likely for the *S. verbesinae* clade. The *H. tuberosus*-associated species *S. longitudinalis* and *S. longipennis* are sister to *H. grosseserratus*-associated species *S. noctipennis* and “Bush’s Fly” respectively. All three of the *Strauzia* SNP phylogenies share topological similarities with the mtDNA phylogeny among the most basal *Strauzia* species.

**Figure 2.**
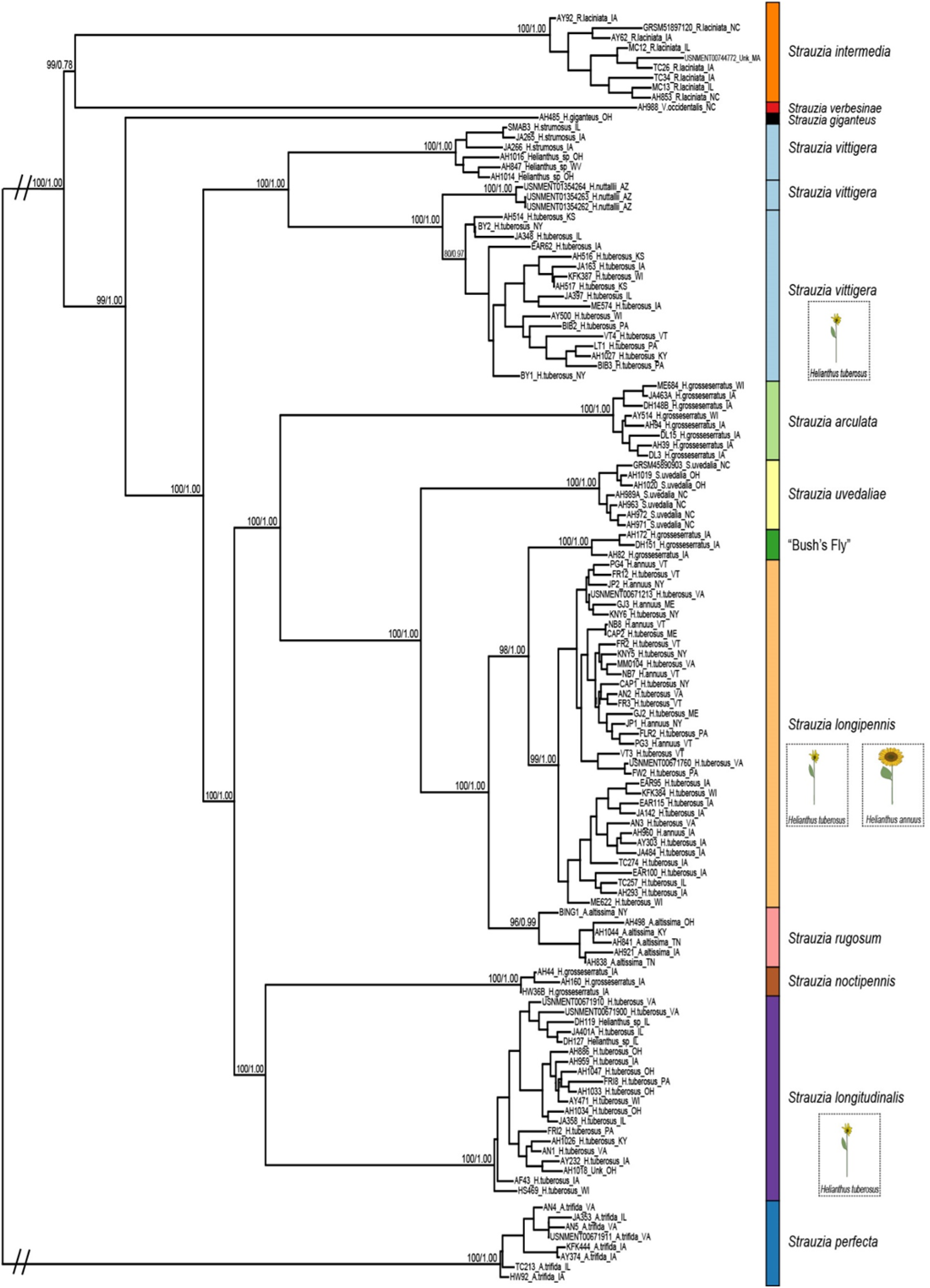
Maximum likelihood phylogeny of *Strauzia* SNP data generated using RAD-sequencing approach. All bootstrap values >75 from the maximum likelihood phylogeny are show on the left, followed by posterior probabilities >0.75 from the Bayesian phylogeny on the right. A general collection site location is indicated after each sample name and specific site locations can be found in Supplemental Table 1. Colored bars for each clade correspond to legend colors in Figure 1. Pictures of *H. tuberosus* are included for all *Strauzia* species collected on that host.

### Phylogenetic discordance and introgression

We explored the possibility of discordant gene tree topologies using a dataset that included one individual of each of 8 *Strauzia* species and a total of 473 loci. The resulting population tree (Figure 3) recapitulates the *Strauzia* topology generated by the Maximum Likelihood and Bayesian SNP phylogenies. The concordance factors at each node range from 0.143 to 0.366 and differ significantly from all alternative topologies that were identified in the analysis. Further, the concordance and population trees generated in BUCKy have the same topology, an indication that the observed gene discordance is the result of incomplete lineage sorting and not introgression. We did not observe any notable differences in the concordance factors or tree topology when varying the α parameter. D-statistics also did not support a significant signature of introgression among the *Strauzia* species sharing the *H. tuberosus* or *H. grosseserratus* hosts, or among closely related species using different hosts (Table 2).

**Table 2.** Results of Patterson’s D-statistics analysis for four *Strauzia* species combinations. *S. perfecta* was the outgroup (O) in all combinations. P3, P2, and P1 represent the three taxa in the analysis, with all analyses testing for introgression between P3 and each of P1 and P2. ABBA represents the number of sites shared between P3 and P2 and BABA represents the number of sites shared between P3 and P1. Equal ABBA and BABA values are expected under the null hypothesis that there is no evidence of introgression and significant deviations from equal ABBA/BABA values provides evidence of introgression among P3 and P2/P1. Statistical significance was evaluated using a Bonferroni correction for multiple tests (α=0.0125). No significant evidence of introgression was detected across any of these four comparisons.

| O | P3 | P2 | P1 | D Statistic | ZScore | ABBA | BABA | Number of Loci | P-Value |
| --- | --- | --- | --- | --- | --- | --- | --- | --- | --- |
| *S. perfecta* | *S. vittigera* | *S. longitudinalis* | *S. longipennis* | -0.05420419 | 0.700739 | 59.74257 | 66.59035 | 490 | 0.48349 |
| *S. perfecta* | *S. uvedaliae* | *S. rugosum* | *S. longipennis* | -0.17811026 | 1.984991 | 26.6593 | 38.21388 | 478 | 0.047156 |
| *S. perfecta* | *S. arcuata* | “Bush’s Fly” | *S. longipennis* | -0.18218319 | 1.393984 | 13.15634 | 19.01795 | 389 | 0.163348 |
| *S. perfecta* | *S. vittigera* | *S. noctipennis* | *S. longitudinalis* | -0.18565298 | 1.884327 | 32.62564 | 47.50148 | 361 | 0.059524 |

**Figure 3.**
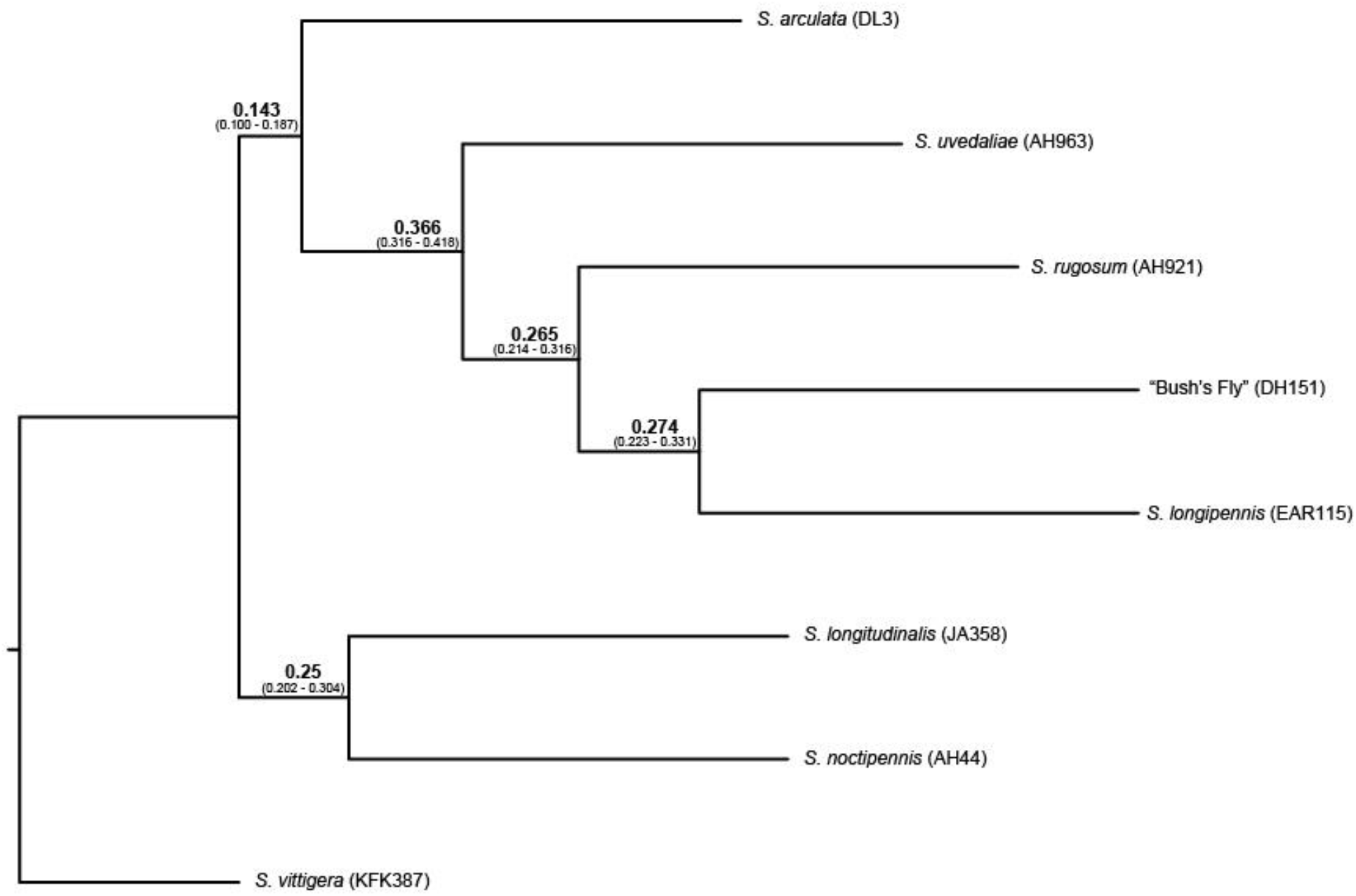
Concordance tree of eight *Strauzia* species generated in BUCKy. Bolded numbers at each node represent the average concordance factor for that node or the average proportion of gene trees that support the topology at the given node. Numbers in parentheses represent the standard deviation of the concordance factor at each node.

## Discussion

### Support for previously named species

This first comprehensive phylogeny of *Strauzia* provides new resolution to several long-standing taxonomic questions. First, all three SNP-based phylogenies support the hypothesis that the genus contains at least 12 reproductively isolated species. Clades on all trees correspond with morphological species concepts proposed by Wiedemann (1830; *S. longipennis*), Lisowski (1985; *S. arculata*, “Bush’s Fly”, *S. intermedia*, *S. longitudinalis*, *S. perfecta*, *S. vittigera*), Steyskal (1986; *S. gigantei*, *S. verbesinae*) and Stoltzfus (1988; *S. noctipennis*, *S. rugosum*, and *S. uvedaliae*). Of all previously named species, only *S. stoltzfusi* (Steyskal 1986) – a species described from a single male / female pair captured on an unknown host in St. Paul, MN – remains in question, and was not available for genetic assessment here. We summarize the outcomes of these results in Table 1 and suggest that a formal morphological re-description of the species in this genus is now warranted.

### *Strauzia* species with multiple hosts – additional host-associated differentiation?

Though most species were reared from a single plant host, two *Strauzia* species were reared from two or more hosts. One of these, S. *vittigera,* contains three distinct clades of flies, each reared from different plant hosts (Figure 2). Two of the *S. vittigera* clades, flies reared from *H. strumosus* and *H. tuberosus,* respectively, come from geographically-overlapping collections, suggesting that the two clades represent reproductively isolated varieties or perhaps distinct species. A third *S. vittigera* clade containing three flies collected from *H. nuttallii* (not reared) in Arizona may also represent a reproductively isolated group, but this requires additional study, as we cannot rule out that their genetic differences may be due to geographic isolation alone. In general, it seems most likely that the morphological species *S. vittigera* is actually a cryptic complex of two or more host-specific species.

*Strauzia longipennis* flies were also reared from two hosts. Flies with this morphology were reared from both *H. tuberosus* and *H. annuus*, but unlike *S. vittigera* these flies showed no apparent host-associated structure. Instead, we resolved two clades of *S. longipennis* that primarily differ in their geographic origin (Midwestern vs. Eastern states). *Strauzia longipennis* may thus be unusual among *Strauzia* flies in its interchangeable use of two plant hosts, with no apparent reproductive isolation having evolved between flies using those hosts. *Helianthus annuus* is the only annual plant used as a host by flies in this genus, so potentially its ubiquity across the landscape, and/or a relaxation of its chemical defenses due to its domestication make it an acceptable spillover host (and see further discussion below).

### Origins of host sharing by host shifting

The primary motivation for this work was determining whether and how multiple closely related insect species come to specialize on the same tissue of the same host. To this end, we find two cases where flies have converged upon the same host, apparently by shifting from other plants. In the first of these cases, *S. arculata*, *S. noctipennis*, and “Bush’s Fly” all feed on the pith of *H. grosseserratus*. If these flies had diverged without host shifting (e.g., via successive periods of geographic isolation and secondary contact), we would expect a phylogeny to show them as one-another’s closest relatives. Instead, each of the three species sharing *H. grosseserratus* as a host is more closely related to a *Strauzia* species on a different host plant (Figure 2).

As with the flies associated with *H. grosseserratus*, the three *Strauzia* species that feed on pith of *H. tuberosus* are not one another’s closest relatives (Figure 2), such that the most parsimonious explanation for the origin of their host sharing is multiple independent shifts onto the same host plant. While previous work using mitochondrial data and microsatellites hypothesized that the *H. tuberosus*-associated flies might have resulted from several recent divergence events during periods of isolation in glacial refugia (Hippee et al. 2016), this was based on the assumption from taxonomic work that the *H. tuberosus* flies were three partially isolated, but otherwise closely related “varieties” of *S. longipennis* (Loew 1873; Foote et al. 1993). The inclusion of other *Strauzia* species in the current work reveals that the *H. tuberosus*-associated flies are actually not one another’s closest relatives, and that they represent distinct, already-isolated lineages that converged upon the same plant.

However, recent phylogenomic study of *H. tuberosus* suggests a more complex origin story for two of its associated *Strauzia* flies. *Helianthus tuberosus* is a hexaploid hybrid of two other sunflowers – *H. grosseserratus* and *H. strumosus* (Bock et al 2014). The *Strauzia* species tree (Figure 2) shows that the *Strauzia longitudinalis* and *Strauzia vittigera* found on *H. tuberosus* both have sister clades of flies that use the *H. grosseserratus* and *H. strumosus* hosts, respectively. One possibility, then, is that these flies both moved onto *H. tuberosus* at the time of its hybrid origin. This scenario does, in one sense, describe the origins of host sharing via two parallel host shifts, but in another it can be viewed as introgression between hosts leading to a convergence of those hosts’ parasites. A future goal will be to collect additional samples and loci to infer divergence dates for these splits – and potentially date the origin of *H. tuberosus*.

### A recent radiation of flies and hosts?

Many *Strauzia* species’ shifts onto the same plant hosts appear to be relatively recent. Incongruence between morphological species and mtDNA data (Axen et al. 2011; Forbes et al. 2013; Figure S2) as well as our gene discordance analysis (Figure 3) and Patterson’s D-statistics calculations (Table 2) all suggest that the radiation of all of the *Strauzia* species associated with *Helianthus* has occurred recently. While we have not yet inferred the divergence timing of *Strauzia* species, recent work in *Helianthus* suggests some intriguing hypotheses. Other than *H. annuus*, all of the *Helianthus* hosts of *Strauzia* belong to a young (1-2 MYA; Mason 2018) clade of perennial sunflowers characterized by rampant interspecific introgression and several incidents of polyploid hybrid speciation (Bock et al. 2014; Baute et al. 2016). The radiation of the perennial sunflowers may have represented an expansion of potential host environments for *Strauzia* and caused a radiation-in-parallel among the flies. Host sharing in this case may be a function of both the recency of origin of some *Helianthus* species and their ongoing sharing of genetic material, which may reduce physiological challenges for flies sampling among plant species.

If host sharing is of recent origin, will it persist? Host sharing might occur temporarily during host shift speciation events in specialist insects, but over longer time periods one species might still “win”. Previous work in *Strauzia* does not support this prediction, instead providing evidence of niche partitioning and prezygotic reproductive barriers that should serve to stabilize host sharing. Primarily, differences in adult emergence timing, combined with short (~18 day) adult life spans, reduce temporal overlap among *Strauzia* species (Hippee et al. 2016). Similar temporal isolation helps maintain species boundaries between congeneric ectoparasites of fish (Šimková and Morand 2008) and sister species of elm leaf beetle that feed on the same host trees in sympatry (Zhang et al. 2015). Premating sexual isolation is also strong in *Strauzia*. Even when males and females of two species are present, conspecific couplings are far more common than interspecific couplings in both the lab (personal observation, ACH) and the field (Hippee et al. 2016). Further, *Strauzia* males that live on shared hosts tend to have more extreme morphological differences from one another than males that do not share hosts (A.H., personal observation), indicating that these traits may be used in discrimination among potential mates and help reduce intersexual conflict.

### Coda: A role for sunflower domestication in a recent *Strauzia* host-shift speciation event?

Our data suggest one additional question for future study: has domestication of sunflowers affected recent *Strauzia* evolution? *Helianthus tuberosus* (Jerusalem artichoke) and *H. annuus* (Common sunflower), the shared hosts of *S. longipennis*, were both grown in the Eastern Agricultural Complex (1-3 KYA) by Native Americans (Delcourt & Delcourt 2004; Kays and Nottingham 2008; Blackman et al. 2011). *Strauzia longipennis* and its sister fly “Bush’s Fly” differ by the shortest genetic distance of all species pairs on the *Strauzia* tree, suggesting a very recent origin. We suggest that the origin of *S. longipennis* may correspond with a shift onto these two sunflower host plants subsequent to their domestication. Future work will employ population genomic and demographic analyses to better understand the impacts of plant domestication on specialist insect speciation and diversification.

## Supporting information

Supplemental Table 2 Page 1

Supplemental Table 2 Page 2

## Acknowledgements

This research required extensive support planning and executing collections trips, and we acknowledge the following for their help: Jarod Armenta, Darius Ballard, Chris Bedel at Edge of Appalachia Nature Preserve, Elana Becker, Stewart Berlocher, Andrew Berry at Bernheim Arboretum and Research Forest, Maggie Blackledge, Charles Bray at F.W. Kent State Park, Zurijanne Carter at Greater Parks of Hamilton County, Sara Childs and other staff at Duke Forest, Alex Cooper, Neisha Croffitt, Sarah Duhon, Maren Elnes, Kristin Fernandez-Kong, Gunther Hansen, Ron Hoham, Demaceo Howard, Jamien and Gregg Jacobs, Patrick Kelly, Dacia Lipkea, Michael Lopez, Jennifer Megyesi at Fat Rooster Farm, Kara Middleton, Erika Mitchell, Kevin Neely, Tom Powell, Emily Reasoner, Halee Schomberg, Bob Smith, Heather Widmayer, Kyle Woods, Pete Woods, and Alex Young. We also acknowledge lab crayfish Prawnsalind Franklin for coral support. Harold Robinson (Smithsonian Institution) helped with the identification of some plant hosts. Funding for the project was provided by grants from the Iowa Academy of Sciences (AAF and ACH), and a CGRER travel grant (ACH). Mention of trade names or commercial products in this publication is solely for the purpose of providing specific information and does not imply recommendation or endorsement by the USDA. USDA is an equal opportunity provider and employer.

## Supplemental Information

- Supplemental Figure 1. Map of *Strauzia* collections in Eastern Iowa including samples used in mtDNA and SNP phylogenetic analyses.
- Supplemental Figure 2. Bayesian phylogeny of *Strauzia* mtCOI gene.
- Supplemental Figure 3. Maximum likelihood phylogeny of 7 *Strauzia* species and *Z. electa*.
- Supplemental Figure 4. Quartet-based phylogenetic analysis of *Strauzia* SNP data generated using SVDQuartets.
- Supplemental Table 1. Table of all collection and DNA extraction information for each specimen included in this manuscript.
- Supplemental Table 2. Table of iTru7 Illumina multiplexing indexes and barcodes used in 3RAD library preparation.

