## Supplemental Table 1 for "Host shifting and host sharing in a genus of specialist flies diversifying alongside their sunflower hosts"

| Sample ID | Species | Dataset | Extraction Method | Latitude | Longitude | Altitude (meters) | State | Life Stage | Sex | Host Plant Species |
|---|---|---|---|---|---|---|---|---|---|---|
| AF43 | *S. longitudinalis* | 3RAD | CTAB | 41.6508 | -91.5602 | 233 | Iowa | Adult | M | *H. tuberosus* |
| AH10 | *S. longipennis* | COI | KIT | 41.4216 | -95.8543 | 377 | Iowa | Adult | M | *H. tuberosus* |
| AH1014 | *S. vittigera* | BOTH | CTAB | 38.7593 | -83.4068 | 226 | Ohio | Larva | N/A | Unknown *Helianthus* |
| AH1016 | *S. vittigera* | BOTH | CTAB | 38.7593 | -83.4068 | 226 | Ohio | Larva | N/A | Unknown *Helianthus* |
| AH1018 | *S. uvedaliae* | 3RAD | CTAB | 38.7593 | -83.4068 | 226 | Ohio | Larva | N/A | *S. uvedalia* |
| AH1019 | *S. uvedaliae* | 3RAD | CTAB | 38.7593 | -83.4068 | 226 | Ohio | Larva | N/A | *S. uvedalia* |
| AH1020 | *S. uvedaliae* | 3RAD | CTAB | 38.7593 | -83.4068 | 226 | Ohio | Larva | N/A | *S. uvedalia* |
| AH1026 | *S. longitudinalis* | 3RAD | CTAB | 38.2109 | -85.4794 | 176 | Kentucky | Larva | N/A | *H. tuberosus* |
| AH1027 | *S. vittigera* | 3RAD | CTAB | 38.2109 | -85.4794 | 176 | Kentucky | Larva | N/A | *H. tuberosus* |
| AH1033 | *S. longitudinalis* | 3RAD | CTAB | 39.1481 | -84.3294 | 152 | Ohio | Larva | N/A | *H. tuberosus* |
| AH1034 | *S. longitudinalis* | 3RAD | CTAB | 39.1481 | -84.3294 | 152 | Ohio | Larva | N/A | *H. tuberosus* |
| AH1044 | *S. rugosum* | 3RAD | CTAB | 37.9159 | -85.6558 | 164 | Kentucky | Larva | N/A | *A. altissima* |
| AH1047 | *S. longitudinalis* | 3RAD | CTAB | 39.2645 | -84.5377 | 215 | Ohio | Larva | N/A | *H. tuberosus* |
| AH160 | *S. noctipennis* | 3RAD | CTAB | 43.4011 | -95.202 | 449 | Iowa | Adult | F | *H. grosseserratus* |
| AH161 | *S. arculata* | COI | KIT | 43.4011 | -95.202 | 449 | Iowa | Adult | F | *H. grosseserratus* |
| AH17 | *S. arculata* | COI | KIT | 43.4011 | -95.202 | 449 | Iowa | Adult | M | *H. grosseserratus* |
| AH172 | "Bush's Fly" | 3RAD | CTAB | 43.4011 | -95.202 | 449 | Iowa | Adult | F | *H. grosseserratus* |
| AH293 | *S. longipennis* | 3RAD | CTAB | 41.6529 | -91.4804 | 217 | Iowa | Adult | M | *H. tuberosus* |
| AH35 | *S. arculata* | COI | KIT | 43.4011 | -95.202 | 449 | Iowa | Adult | M | *H. grosseserratus* |
| AH39 | *S. arculata* | 3RAD | CTAB | 43.4011 | -95.202 | 449 | Iowa | Adult | M | *H. grosseserratus* |
| AH44 | *S. noctipennis* | 3RAD | CTAB | 43.4011 | -95.202 | 449 | Iowa | Adult | M | *H. grosseserratus* |
| AH45 | *S. noctipennis* | COI | KIT | 43.4011 | -95.202 | 449 | Iowa | Adult | M | *H. grosseserratus* |
| AH471 | Unknown | COI | CTAB | 41.1244 | -81.5179 | 236 | Ohio | Larva | N/A | *H. tuberosus* |
| AH472 | *S. rugosum* | COI | CTAB | 39.5892 | -82.5744 | 234 | Ohio | Larva | N/A | *A. altissima* |
| AH473 | Unknown | COI | CTAB | 41.1244 | -81.5179 | 236 | Ohio | Larva | N/A | *H. tuberosus* |
| AH474 | Unknown | COI | CTAB | 41.7048 | -85.0215 | 297 | Indiana | Larva | N/A | *H. tuberosus* |
| AH481 | *S. gigantei* | COI | CTAB | 41.5909 | -83.7616 | 197 | Ohio | Larva | N/A | *H. giganteus* |
| AH483 | *S. rugosum* | COI | CTAB | 38.9484 | -85.7968 | 172 | Indiana | Larva | N/A | *A. altissima* |
| AH485 | *S. gigantei* | 3RAD | CTAB | 41.5909 | -83.7616 | 198 | Ohio | Larva | N/A | *H. giganteus* |
| AH488 | *S. gigantei* | COI | CTAB | 41.5909 | -83.7616 | 198 | Ohio | Larva | N/A | *H. giganteus* |
| AH494 | *S. rugosum* | COI | CTAB | 41.1244 | -81.5179 | 236 | Ohio | Larva | N/A | *A. altissima* |
| AH498 | *S. rugosum* | 3RAD | CTAB | 41.1244 | -81.5179 | 236 | Ohio | Larva | N/A | *A. altissima* |
| AH500 | Unknown | COI | CTAB | 39.5892 | -82.5744 | 234 | Ohio | Larva | N/A | Unknown *Helianthus* |
| AH501 | Unknown | COI | CTAB | 39.5892 | -82.5744 | 234 | Ohio | Larva | N/A | Unknown *Helianthus* |
| AH514 | *S. vittigera* | BOTH | CTAB | 39.1471 | -96.449 | 312 | Kansas | larva | N/A | *H. tuberosus* |
| AH516 | *S. vittigera* | BOTH | CTAB | 39.1471 | -96.449 | 312 | Kansas | larva | N/A | *H. tuberosus* |
| AH517 | *S. vittigera* | BOTH | CTAB | 39.1471 | -96.449 | 312 | Kansas | larva | N/A | *H. tuberosus* |
| AH519 | Unknown | COI | CTAB | 39.1471 | -96.449 | 312 | Kansas | larva | N/A | *H. tuberosus* |
| AH522 | *S. rugosum* | COI | CTAB | 39.1471 | -96.449 | 312 | Kansas | larva | N/A | *A. altissima* |
| AH523 | *S. rugosum* | COI | CTAB | 39.1471 | -96.449 | 312 | Kansas | larva | N/A | *A. altissima* |
| AH527 | Unknown | COI | CTAB | 39.1471 | -96.449 | 312 | Kansas | larva | N/A | *H. tuberosus* |
| AH55 | *S. arculata* | COI | KIT | 43.4011 | -95.202 | 449 | Iowa | Adult | M | *H. grosseserratus* |
| AH82 | "Bush's Fly" | 3RAD | CTAB | 43.401 | -95.2498 | 433 | Iowa | Adult | M | *H. grosseserratus* |
| AH838 | *S. rugosum* | 3RAD | CTAB | 36.9855 | -88.2063 | 130 | Tennessee | Larva | N/A | *A. altissima* |
| AH841 | *S. rugosum* | 3RAD | CTAB | 36.9855 | -88.2063 | 130 | Tennessee | Larva | N/A | *A. altissima* |
| AH847 | *S. vittigera* | 3RAD | CTAB | 38.2732 | -81.6588 | 231 | West Virginia | Larva | N/A | Unknown *Helianthus* |
| AH853 | *S. intermedia* | 3RAD | CTAB | 39.8581 | -84.1777 | 258 | Ohio | Larva | N/A | *R. laciniata* |
| AH88 | *S. noctipennis* | COI | KIT | 43.401 | -95.2498 | 433 | Iowa | Adult | M | *H. grosseserratus* |
| AH886 | *S. longitudinalis* | 3RAD | CTAB | 39.8581 | -84.1777 | 258 | Ohio | Larva | N/A | *H. tuberosus* |
| AH921 | *S. rugosum* | 3RAD | CTAB | 41.6652 | -91.5134 | 208 | Iowa | Larva | N/A | *A. altissima* |
| AH94 | *S. arculata* | BOTH | CTAB | 43.4011 | -95.202 | 449 | Iowa | Adult | M | *H. grosseserratus* |
| AH959 | *S. longitudinalis* | 3RAD | CTAB | 41.4788 | -93.9109 | 302 | Iowa | Adult | M | *H. tuberosus* |
| AH960 | *S. longipennis* | 3RAD | CTAB | 41.4788 | -93.9109 | 302 | Iowa | Adult | F | *H. annuus* |
| AH963 | *S. uvedaliae* | 3RAD | CTAB | 36.1599 | -79.4887 | 184 | North Carolina | Larva | N/A | *S. uvedalia* |
| AH971 | *S. uvedaliae* | 3RAD | CTAB | 35.9243 | -79.0502 | 88 | North Carolina | Adult | M | *S. uvedalia* |
| AH972 | *S. uvedaliae* | 3RAD | CTAB | 35.9243 | -79.0502 | 88 | North Carolina | Adult | M | *S. uvedalia* |
| AH977 | *S. verbesina* | COI | CTAB | 36.0112 | -79.0099 | 123 | North Carolina | Larva | N/A | *V. occidentalis* |
| AH988 | *S. verbesina* | BOTH | CTAB | 35.8168 | -81.1785 | 268 | North Carolina | Larva | N/A | *V. occidentalis* |
| AH989A | *S. uvedaliae* | 3RAD | CTAB | 35.8168 | -81.1785 | 268 | North Carolina | Larva | N/A | *S. uvedalia* |
| AN1 | *S. longitudinalis* | 3RAD | CTAB | 38.8828 | -77.1489 | 73 | Virginia | Adult | M | *H. tuberosus* |
| AN2 | *S. longipennis* | BOTH | CTAB | 38.8897 | -77.1419 | 73 | Virginia | Larva | N/A | *H. tuberosus* |
| AN3 | *S. longipennis* | 3RAD | CTAB | 38.8897 | -77.1419 | 73 | Virginia | Larva | N/A | *H. tuberosus* |
| AN4 | *S. perfecta* | 3RAD | CTAB | 38.8631 | -77.1193 | 73 | Virginia | Larva | N/A | *A. trifida* |
| AN5 | *S. perfecta* | 3RAD | CTAB | 38.8631 | -77.1193 | 73 | Virginia | Larva | N/A | *A. trifida* |
| AY232 | *S. longitudinalis* | 3RAD | CTAB | 41.9245 | -91.4167 | 258 | Iowa | Adult | M | *H. tuberosus* |
| AY303 | *S. longipennis* | 3RAD | CTAB | 41.6529 | -91.4804 | 217 | Iowa | Adult | F | *H. tuberosus* |
| AY374 | *S. perfecta* | 3RAD | CTAB | 41.9293 | -91.4235 | 258 | Iowa | Adult | M | *A. trifida* |
| AY471 | *S. longitudinalis* | 3RAD | CTAB | 42.884 | -89.8572 | 311 | Wisconsin | Adult | M | *H. tuberosus* |
| AY500 | *S. vittigera* | 3RAD | CTAB | 42.884 | -89.8572 | 311 | Wisconsin | Adult | M | *H. tuberosus* |
| AY514 | *S. arculata* | 3RAD | CTAB | 43.2975 | -90.3013 | 219 | Wisconsin | Adult | M | *H. grosseserratus* |

| Sample ID | Species | Dataset | Extraction Method | Latitude | Longitude | Altitude (meters) | State | Life Stage | Sex | Host Plant Species |
|---|---|---|---|---|---|---|---|---|---|---|
| AY521 | *S. arculata* | COI | KIT | 43.2975 | -90.3013 | 219 | Wisconsin | Adult | M | *H. grosseserratus* |
| AY62 | *S. intermedia* | 3RAD | CTAB | 43.3012 | -91.7908 | 269 | Iowa | Adult | M | *R. laciniata* |
| AY92 | *S. intermedia* | 3RAD | CTAB | 43.3012 | -91.7908 | 269 | Iowa | Adult | M | *R. laciniata* |
| BIB2 | *S. vittigera* | 3RAD | CTAB | 40.6053 | -79.3706 | 295 | Pennsylvania | Larva | N/A | *H. tuberosus* |
| BIB3 | *S. vittigera* | 3RAD | CTAB | 40.6053 | -79.3706 | 295 | Pennsylvania | Larva | N/A | *H. tuberosus* |
| BING1 | *S. rugosum* | 3RAD | CTAB | 42.0918 | -75.961 | 264 | New York | Larva | N/A | *A. altissima* |
| BY1 | *S. vittigera* | 3RAD | CTAB | 42.7725 | -75.4928 | 366 | New York | Larva | N/A | *H. tuberosus* |
| BY2 | *S. vittigera* | 3RAD | CTAB | 42.7725 | -74.4928 | 366 | New York | Larva | N/A | *H. tuberosus* |
| CAP1 | *S. longipennis* | 3RAD | CTAB | 43.6673 | -70.307 | 12 | Maine | Adult | M | *H. tuberosus* |
| CAP2 | *S. longipennis* | 3RAD | CTAB | 43.6673 | -70.307 | 12 | Maine | Adult | M | *H. tuberosus* |
| DB2 | *S. longipennis* | COI | KIT | 41.9245 | -91.4167 | 258 | Iowa | Adult | M | *H. tuberosus* |
| DH116 | *S. vittigera* | COI | CTAB | 40.1131 | -88.1587 | 214 | Illinois | Adult | M | Unknown *Helianthus* |
| DH119 | *S. vittigera* | 3RAD | CTAB | 40.1131 | -88.1587 | 214 | Illinois | Adult | M | Unknown *Helianthus* |
| DH126A | *S. perfecta* | COI | KIT | 40.1131 | -88.1587 | 214 | Illinois | Adult | F | *A. trifida* |
| DH127 | *S. longitudinalis* | 3RAD | CTAB | 40.1131 | -88.1587 | 214 | Illinois | Adult | M | Unknown *Helianthus* |
| DH148B | "Bush's Fly" | 3RAD | CTAB | 41.0131 | -94.9548 | 371 | Iowa | Adult | M | *H. grosseserratus* |
| DH151 | *S. arculata* | 3RAD | CTAB | 41.0131 | -94.9548 | 371 | Iowa | Adult | M | *H. grosseserratus* |
| DH152 | *S. arculata* | COI | KIT | 41.0131 | -94.9548 | 371 | Iowa | Adult | M | *H. grosseserratus* |
| DH159 | *S. noctipennis* | COI | KIT | 41.0131 | -94.9548 | 371 | Iowa | Adult | M | *H. grosseserratus* |
| DH79 | *S. intermedia* | COI | KIT | 40.197 | -88.3913 | 205 | Illinois | Adult | F | *R. laciniata* |
| DL15 | *S. arculata* | 3RAD | CTAB | 41.7355 | -91.7225 | 257 | Iowa | Adult | M | *H. grosseserratus* |
| DL3 | *S. arculata* | 3RAD | CTAB | 41.7355 | -91.7225 | 257 | Iowa | Adult | M | *H. grosseserratus* |
| EAR100 | *S. longipennis* | 3RAD | CTAB | 41.693 | -91.548 | 203 | Iowa | Adult | M | *H. tuberosus* |
| EAR115 | *S. longipennis* | 3RAD | CTAB | 41.6958 | -92.2797 | 263 | Iowa | Adult | F | *H. tuberosus* |
| EAR62 | *S. vittigera* | 3RAD | CTAB | 41.6529 | -91.4804 | 217 | Iowa | Adult | M | *H. tuberosus* |
| EAR95 | *S. longipennis* | 3RAD | CTAB | 41.66 | -91.5775 | 209 | Iowa | Adult | M | *H. tuberosus* |
| FLR2 | *S. longipennis* | BOTH | CTAB | 40.1347 | -79.7464 | 246 | Pennsylvania | Larva | N/A | *H. tuberosus* |
| FR12 | *S. longipennis* | 3RAD | CTAB | 43.8532 | -72.589 | 184 | Vermont | Adult | M | *H. tuberosus* |
| FR2 | *S. longipennis* | 3RAD | CTAB | 43.8532 | -72.589 | 184 | Vermont | Adult | M | *H. tuberosus* |
| FR3 | *S. longipennis* | 3RAD | CTAB | 43.8532 | -72.589 | 184 | Vermont | Adult | M | *H. tuberosus* |
| FRI2 | *S. longitudinalis* | 3RAD | CTAB | 40.4454 | -79.9039 | 297 | Pennsylvania | Larva | N/A | *H. tuberosus* |
| FRI8 | *S. longitudinalis* | 3RAD | CTAB | 40.4454 | -79.9039 | 297 | Pennsylvania | Larva | N/A | *H. tuberosus* |
| FW2 | *S. longipennis* | 3RAD | CTAB | 40.1266 | -75.1704 | 60 | Pennsylvania | Adult | M | *H. tuberosus* |
| GBDP18754 | *Paramyiolia rhino* | COI | GENBANK | N/A | N/A | N/A | N/A | N/A | N/A | N/A |
| GBDP18755 | *Paramyiolia rhino* | COI | GENBANK | N/A | N/A | N/A | N/A | N/A | N/A | N/A |
| GBDP18756 | *Paramyiolia rhino* | COI | GENBANK | N/A | N/A | N/A | N/A | N/A | N/A | N/A |
| GJ2 | *S. longipennis* | 3RAD | CTAB | 43.6667 | -70.2921 | 37 | Maine | Adult | M | *R. laciniata* |
| GJ3 | *S. longipennis* | BOTH | CTAB | 43.6667 | -70.2921 | 37 | Maine | Adult | M | *S. uvedalia* |
| GRSM10541862 | *S. rugosum* | COI | KIT | 35.5844 | -83.0728 | 1463 | North Carolina | Adult | N/A | *S. uvedalia* |
| GRSM18565392 | *S. rugosum* | COI | KIT | 35.5844 | -83.0728 | 1463 | North Carolina | Adult | N/A | N/A |
| GRSM27015179 | *S. rugosum* | COI | KIT | 35.5844 | -83.0728 | 1463 | North Carolina | Adult | N/A | N/A |
| GRSM45890903 | *S. uvedaliae* | 3RAD | CTAB | 35.6777 | -83.5371 | 566 | Tennessee | Adult | M | N/A |
| GRSM51897120 | *S. intermedia* | BOTH | CTAB/KIT | 35.5844 | -83.0728 | 1463 | North Carolina | Adult | M | *H. annuus* |
| GRSM55049074 | *S. uvedaliae* | COI | KIT | 35.7033 | -83.3866 | 537 | Tennessee | Adult | N/A | *H. annuus* |
| GRSM71385783 | *S. uvedaliae* | COI | KIT | 35.7033 | -83.3866 | 537 | Tennessee | Adult | M | *S. uvedalia* |
| HH02 | *S. arculata* | COI | KIT | 41.6652 | -91.5134 | 208 | Iowa | Adult | F | *H. grosseserratus* |
| HH04 | *S. arculata* | COI | KIT | 41.6652 | -91.5134 | 208 | Iowa | Adult | F | *H. grosseserratus* |
| HS469 | *S. longitudinalis* | 3RAD | CTAB | 42.884 | -89.8572 | 311 | Wisconsin | Adult | M | *H. tuberosus* |
| HW36B | *S. noctipennis* | 3RAD | CTAB | 41.7355 | -91.7225 | 257 | Iowa | Adult | M | *H. grosseserratus* |
| HW92 | *S. perfecta* | 3RAD | CTAB | 41.6529 | -91.4804 | 217 | Iowa | Adult | M | *A. trifida* |
| JA1 | *S. longitudinalis* | COI | CTAB | 41.9245 | -91.4167 | 258 | Iowa | Adult | M | *H. tuberosus* |
| JA116 | *S. vittigera* | COI | KIT | 41.4216 | -95.8543 | 377 | Iowa | Adult | F | *H. tuberosus* |
| JA135 | *S. longipennis* | COI | KIT | 41.4216 | -95.8543 | 377 | Iowa | Adult | M | *H. tuberosus* |
| JA142 | *S. longipennis* | 3RAD | CTAB | 41.4216 | -95.8543 | 377 | Iowa | Adult | M | *H. tuberosus* |
| JA149 | *S. longipennis* | COI | KIT | 41.4216 | -95.8543 | 377 | Iowa | Adult | F | *H. tuberosus* |
| JA154 | *S. longitudinalis* | COI | KIT | 41.4216 | -95.8543 | 377 | Iowa | Adult | F | *H. tuberosus* |
| JA163 | *S. vittigera* | 3RAD | CTAB | 41.4216 | -95.8543 | 377 | Iowa | Adult | M | *H. tuberosus* |
| JA261 | *S. vittigera* | COI | KIT | 41.9154 | -91.5162 | 241 | Iowa | Adult | F | *H. strumosus* |
| JA265 | *S. vittigera* | 3RAD | CTAB | 41.9154 | -91.5162 | 241 | Iowa | Adult | M | *H. strumosus* |
| JA266 | *S. vittigera* | 3RAD | CTAB | 41.9154 | -91.5162 | 241 | Iowa | Adult | M | *H. strumosus* |
| JA285 | *S. perfecta* | COI | KIT | 40.1128 | -88.1336 | 211 | Illinois | Adult | M | *A. trifida* |
| JA348 | *S. vittigera* | 3RAD | CTAB | 40.1131 | -88.1587 | 214 | Illinois | Adult | M | *H. tuberosus* |
| JA353 | *S. perfecta* | 3RAD | CTAB | 40.1131 | -88.1587 | 214 | Illinois | Adult | M | *A. trifida* |
| JA358 | *S. longitudinalis* | 3RAD | CTAB | 40.1128 | -88.1336 | 211 | Illinois | Adult | M | *H. tuberosus* |
| JA359 | *S. longitudinalis* | COI | KIT | 40.1128 | -88.1336 | 211 | Illinois | Adult | M | *H. tuberosus* |
| JA361 | *S. vittigera* | COI | KIT | 40.1128 | -88.1336 | 211 | Illinois | Adult | F | *H. tuberosus* |
| JA365 | *S. vittigera* | COI | KIT | 40.1128 | -88.1336 | 211 | Illinois | Adult | M | *H. tuberosus* |
| JA381 | *S. longipennis* | COI | CTAB | 40.1128 | -88.1336 | 211 | Illinois | Adult | M | *H. tuberosus* |
| JA397 | *S. vittigera* | 3RAD | CTAB | 40.1131 | -88.1587 | 214 | Illinois | Adult | M | *H. tuberosus* |
| JA401A | *S. longitudinalis* | 3RAD | CTAB | 40.1131 | -88.1587 | 214 | Illinois | Adult | M | *H. tuberosus* |
| JA463A | *S. arculata* | 3RAD | CTAB | 41.0131 | -94.9548 | 371 | Iowa | Adult | M | *H. grosseserratus* |

| Sample ID | Species | Dataset | Extraction Method | Latitude | Longitude | Altitude (meters) | State | Life Stage | Sex | Host Plant Species |
|---|---|---|---|---|---|---|---|---|---|---|
| JA484 | *S. longipennis* | 3RAD | CTAB | 41.4216 | -95.8543 | 377 | Iowa | Adult | M | *H. tuberosus* |
| JA566 | *S. longipennis* | COI | CTAB | 42.884 | -89.8572 | 311 | Wisconsin | Adult | M | *H. tuberosus* |
| JA613 | *S. vittigera* | COI | KIT | 42.884 | -89.8572 | 311 | Wisconsin | Adult | M | *H. tuberosus* |
| JA62 | *S. intermedia* | COI | KIT | 43.3012 | -91.7908 | 269 | Iowa | Adult | M | *R. laciniata* |
| JA636 | *S. longipennis* | COI | KIT | 43.4461 | -90.3632 | 231 | Wisconsin | Adult | M | *H. tuberosus* |
| JP1 | *S. longipennis* | 3RAD | CTAB | 42.7058 | -73.6682 | 85 | New York | Adult | M | *H. annuus* |
| JP2 | *S. longipennis* | 3RAD | CTAB | 42.7058 | -73.6682 | 85 | New York | Adult | M | *H. annuus* |
| KFK384 | *S. longipennis* | 3RAD | CTAB | 42.884 | -89.8572 | 311 | Wisconsin | Adult | M | *H. tuberosus* |
| KFK387 | *S. vittigera* | 3RAD | CTAB | 42.884 | -89.8572 | 311 | Wisconsin | Adult | M | *H. tuberosus* |
| KFK404 | *S. arculata* | COI | KIT | 43.2975 | -90.3013 | 219 | Wisconsin | Adult | M | *H. grosseserratus* |
| KFK444 | *S. perfecta* | BOTH | CTAB | 41.9657 | -91.5797 | 219 | Iowa | Adult | M | *A. trifida* |
| KNY5 | *S. longipennis* | 3RAD | CTAB | 41.2612 | -73.687 | 75 | New York | Adult | M | *H. tuberosus* |
| KNY6 | *S. longipennis* | 3RAD | CTAB | 41.2612 | -73.687 | 75 | New York | Adult | M | *H. tuberosus* |
| LT1 | *S. vittigera* | BOTH | CTAB | 40.4625 | -78.1998 | 257 | Pennsylvania | Larva | N/A | *H. tuberosus* |
| MC12 | *S. intermedia* | 3RAD | CTAB | 40.1974 | -88.3821 | 205 | Illinois | Adult | M | *R. laciniata* |
| MC13 | *S. intermedia* | 3RAD | CTAB | 40.1974 | -88.3821 | 205 | Illinois | Adult | M | *R. laciniata* |
| ME22 | *S. intermedia* | COI | KIT | 43.3012 | -91.7908 | 269 | Iowa | Adult | M | *R. laciniata* |
| ME37 | *S. vittigera* | COI | KIT | 40.1974 | -88.3821 | 205 | Illinois | Adult | M | *H. tuberosus* |
| ME41 | *S. arculata* | COI | KIT | 41.0131 | -94.9548 | 371 | Iowa | Adult | M | *H. grosseserratus* |
| ME51 | *S. longipennis* | COI | KIT | 41.4216 | -95.8543 | 377 | Iowa | Adult | M | *H. tuberosus* |
| ME574 | *S. vittigera* | 3RAD | CTAB | 41.6529 | -91.4804 | 217 | Iowa | Adult | M | *H. tuberosus* |
| ME622 | *S. longipennis* | 3RAD | CTAB | 42.884 | -89.8572 | 311 | Wisconsin | Adult | M | *H. tuberosus* |
| ME684 | *S. arculata* | 3RAD | CTAB | 43.2975 | -90.3013 | 219 | Wisconsin | Adult | M | *H. grosseserratus* |
| MM0104 | *S. longipennis* | 3RAD | CTAB | 38.8828 | -77.1489 | 79 | Virginia | Adult | M | Unknown *Helianthus* |
| NB7 | *S. longipennis* | 3RAD | CTAB | 44.2847 | -72.5749 | 169 | Vermont | Adult | M | *H. annuus* |
| NB8 | *S. longipennis* | 3RAD | CTAB | 44.2847 | -72.5749 | 169 | Vermont | Adult | M | *H. annuus* |
| PG3 | *S. longipennis* | BOTH | CTAB | 41.9045 | -71.9038 | 67 | Connecticut | Adult | M | *H. annuus* |
| PG4 | *S. longipennis* | 3RAD | CTAB | 41.9045 | -71.9038 | 67 | Connecticut | Adult | M | *H. annuus* |
| SMAB3 | *S. vittigera* | 3RAD | CTAB | 41.914 | -88.36 | 225 | Illinois | Adult | M | *H. strumosus* |
| st0015 | *S. longitudinalis* | COI | KIT | 41.9657 | -91.5797 | 219 | Iowa | Adult | F | *H. tuberosus* |
| st0026 | *S. vittigera* | COI | KIT | 41.9657 | -91.5797 | 219 | Iowa | Adult | F | *H. tuberosus* |
| st0032 | *S. vittigera* | COI | KIT | 41.6477 | -91.5703 | 208 | Iowa | Adult | M | *H. tuberosus* |
| st0037 | *S. longitudinalis* | COI | KIT | 41.9245 | -91.4167 | 258 | Iowa | Adult | F | *H. tuberosus* |
| st0086 | *S. longipennis* | COI | KIT | 41.9657 | -91.5797 | 219 | Iowa | Adult | M | *H. tuberosus* |
| st0097 | *S. longipennis* | COI | KIT | 41.9657 | -91.5797 | 219 | Iowa | Adult | F | *H. tuberosus* |
| ST058 | *S. longipennis* | COI | KIT | 41.6477 | -91.5703 | 208 | Iowa | Adult | M | *H. tuberosus* |
| st077 | *S. longitudinalis* | COI | KIT | 41.9245 | -91.4167 | 258 | Iowa | Adult | M | *H. tuberosus* |
| STTC7 | *S. noctipennis* | COI | KIT | 41.747 | -91.5154 | 220 | Iowa | Adult | M | *H. grosseserratus* |
| TC15 | *S. arculata* | COI | KIT | 41.747 | -90.5154 | 220 | Iowa | Adult | M | *H. grosseserratus* |
| TC2 | *S. noctipennis* | COI | KIT | 41.747 | -89.5154 | 220 | Iowa | Adult | M | *H. grosseserratus* |
| TC213 | *S. perfecta* | 3RAD | CTAB | 40.1131 | -88.1587 | 214 | Illinois | Adult | F | *A. trifida* |
| tc257 | *S. longipennis* | 3RAD | CTAB | 40.1131 | -88.1587 | 214 | Illinois | Adult | F | *H. tuberosus* |
| TC26 | *S. intermedia* | 3RAD | CTAB | 41.6657 | -91.5127 | 208 | Iowa | Adult | M | *R. laciniata* |
| TC274 | *S. longipennis* | 3RAD | CTAB | 40.1128 | -88.1336 | 211 | Illinois | Adult | M | *H. tuberosus* |
| TC315 | *S. longipennis* | COI | CTAB | 40.1128 | -88.1336 | 211 | Illinois | Adult | M | *H. tuberosus* |
| TC34 | *S. intermedia* | 3RAD | CTAB | 43.3012 | -91.7908 | 269 | Iowa | Adult | M | *R. laciniata* |
| TC373 | *S. longipennis* | COI | KIT | 41.6529 | -91.4804 | 217 | Iowa | Adult | F | *H. tuberosus* |
| TC75 | *S. intermedia* | COI | KIT | 43.3012 | -91.7908 | 269 | Iowa | Adult | F | *R. laciniata* |
| TC81 | *S. intermedia* | COI | KIT | 40.1974 | -88.3821 | 205 | Illinois | Adult | M | *R. laciniata* |
| USNMENT00671213 | *S. longipennis* | 3RAD | CTAB | 38.8722 | -77.1322 | 67 | Virginia | Adult | F | *H. tuberosus* |
| USNMENT00671214 | *S. longitudinalis* | COI | KIT | 38.8722 | -77.1322 | 67 | Virginia | Adult | M | *H. tuberosus* |
| USNMENT00671760 | *S. longipennis* | 3RAD | CTAB | 38.8783 | -77.1386 | 73 | Virginia | Adult | M | *H. tuberosus* |
| USNMENT00671900 | *S. longitudinalis* | 3RAD | CTAB | 38.8783 | -77.1386 | 73 | Virginia | Adult | M | *H. grosseserratus* |
| USNMENT00671910 | *S. longitudinalis* | 3RAD | CTAB | 38.8792 | -77.1386 | 73 | Virginia | Adult | M | *H. tuberosus* |
| USNMENT00671911 | *S. perfecta* | BOTH | CTAB | 38.8828 | -77.1489 | 79 | Virginia | Adult | F | *H. tuberosus* |
| USNMENT00671914 | *Chetostoma rubidium* | COI | KIT | 35.9172 | -106.5916 | 2625 | New Mexico | Adult | N/A | *H. tuberosus* |
| USNMENT00744772 | *S. intermedia* | 3RAD | CTAB | 42.3629 | -72.4598 | 202 | Massachussetts | Adult | M | *H. tuberosus* |
| USNMENT00745565 | *Trypeta flaveola* | COI | KIT | 44.5042 | -109.9692 | 2140 | Wyoming | Adult | N/A | *A. trifida* |
| USNMENT00745589 | *S. arculata* | COI | KIT | 40.7033 | -104.9986 | 1585 | Colorado | Adult | F | Unknown |
| USNMENT01354262 | *S. vittigera* | 3RAD | CTAB | 34 | -109.4579 | 2560 | Arizona | Adult | M | *H. nuttallii* (not reared) |
| USNMENT01354263 | *S. vittigera* | 3RAD | CTAB | 34 | -109.4579 | 2560 | Arizona | Adult | M | *H. nuttallii* (not reared) |
| USNMENT01354264 | *S. vittigera* | 3RAD | CTAB | 34 | -109.4579 | 2560 | Arizona | Adult | F | *H. nuttallii* (not reared) |
| VT3 | *S. longipennis* | 3RAD | CTAB | 44.2569 | -72.5171 | 245 | Vermont | Adult | M | *H. tuberosus* |
| VT4 | *S. vittigera* | 3RAD | CTAB | 44.2569 | -72.5171 | 245 | Vermont | Adult | M | *H. tuberosus* |
| ZONO1 | *Z. electa* | 3RAD | CTAB | 41.89 | -91.6004 | 234 | Iowa | Adult | M | *Solanum* species |
| ZONO2 | *Z. electa* | 3RAD | CTAB | 41.89 | -91.6004 | 234 | Iowa | Adult | F | *Solanum* species |
