## Supplementary figures and images for "Host shifting and host sharing in a genus of specialist flies diversifying alongside their sunflower hosts"

### Supplemental Figure 1

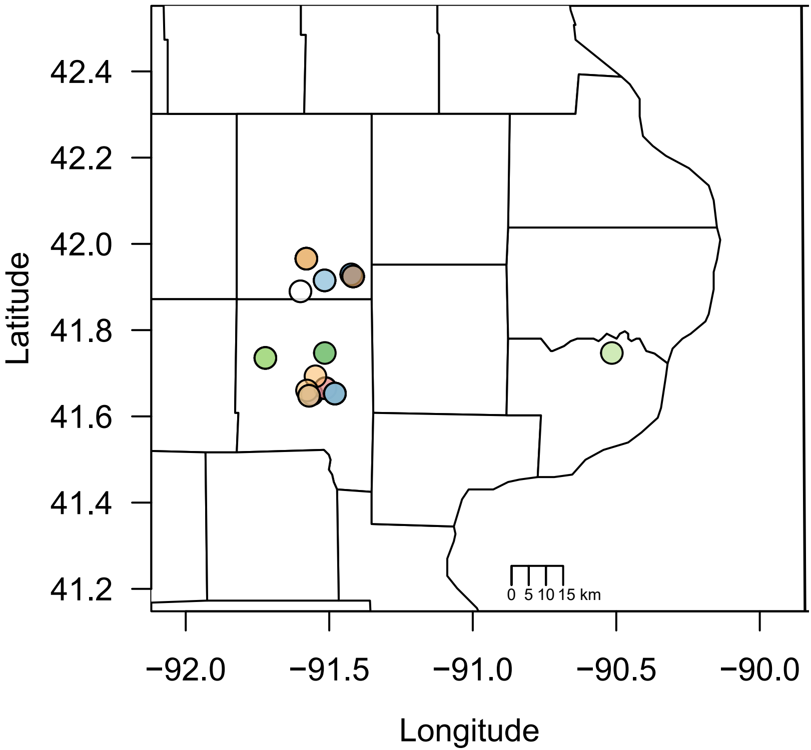

### Supplemental Figure 2

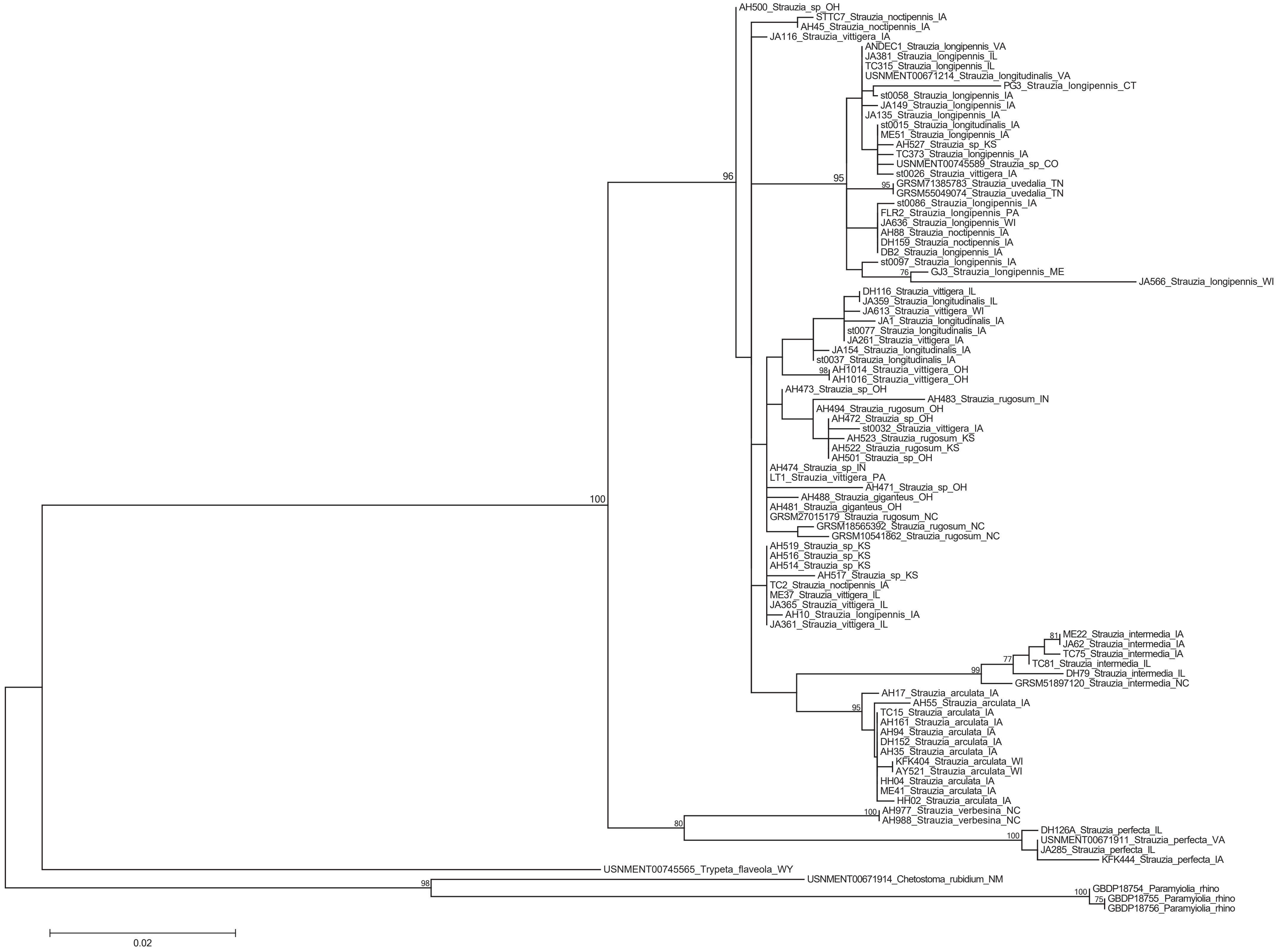

### Supplemental Figure 3

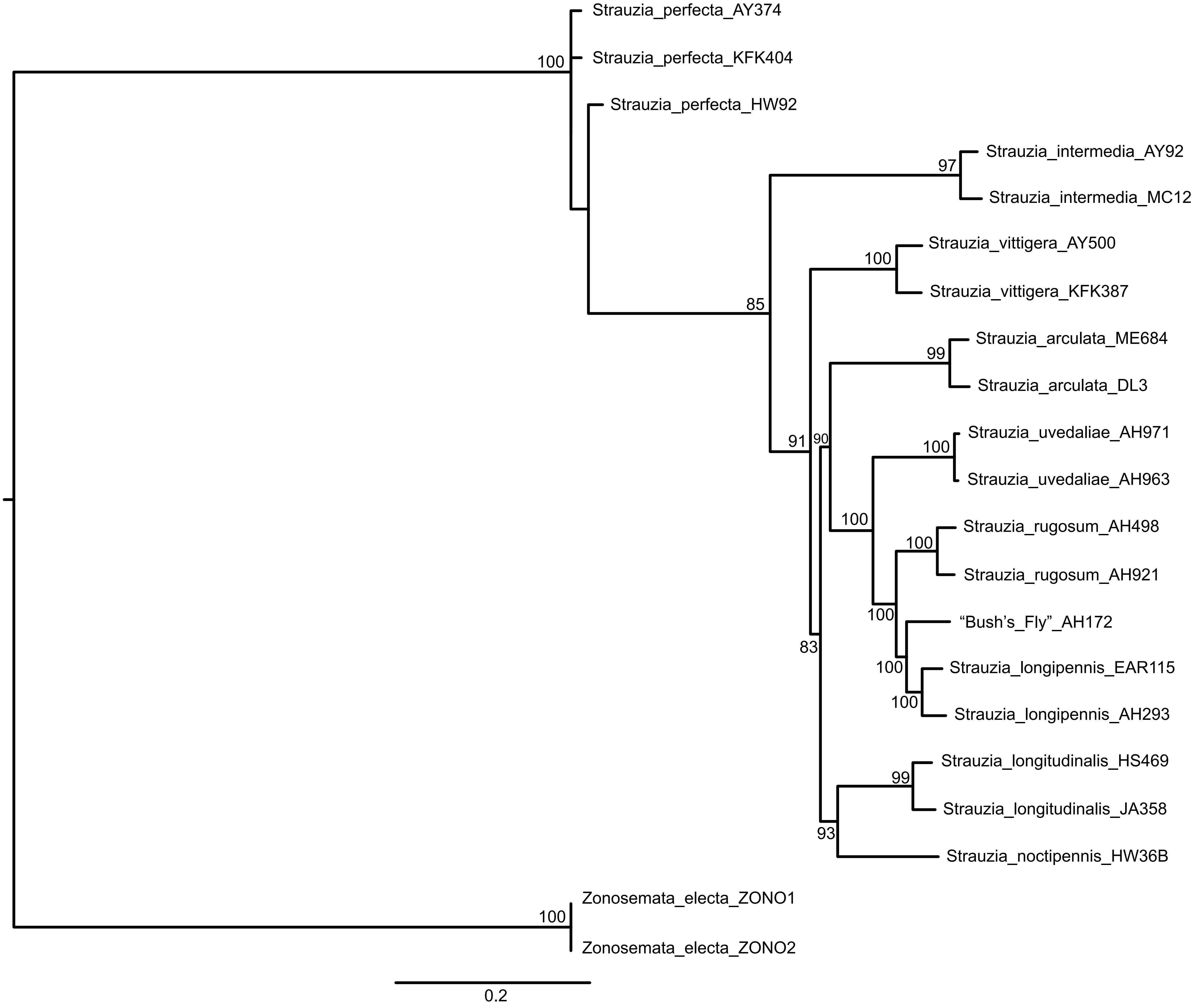

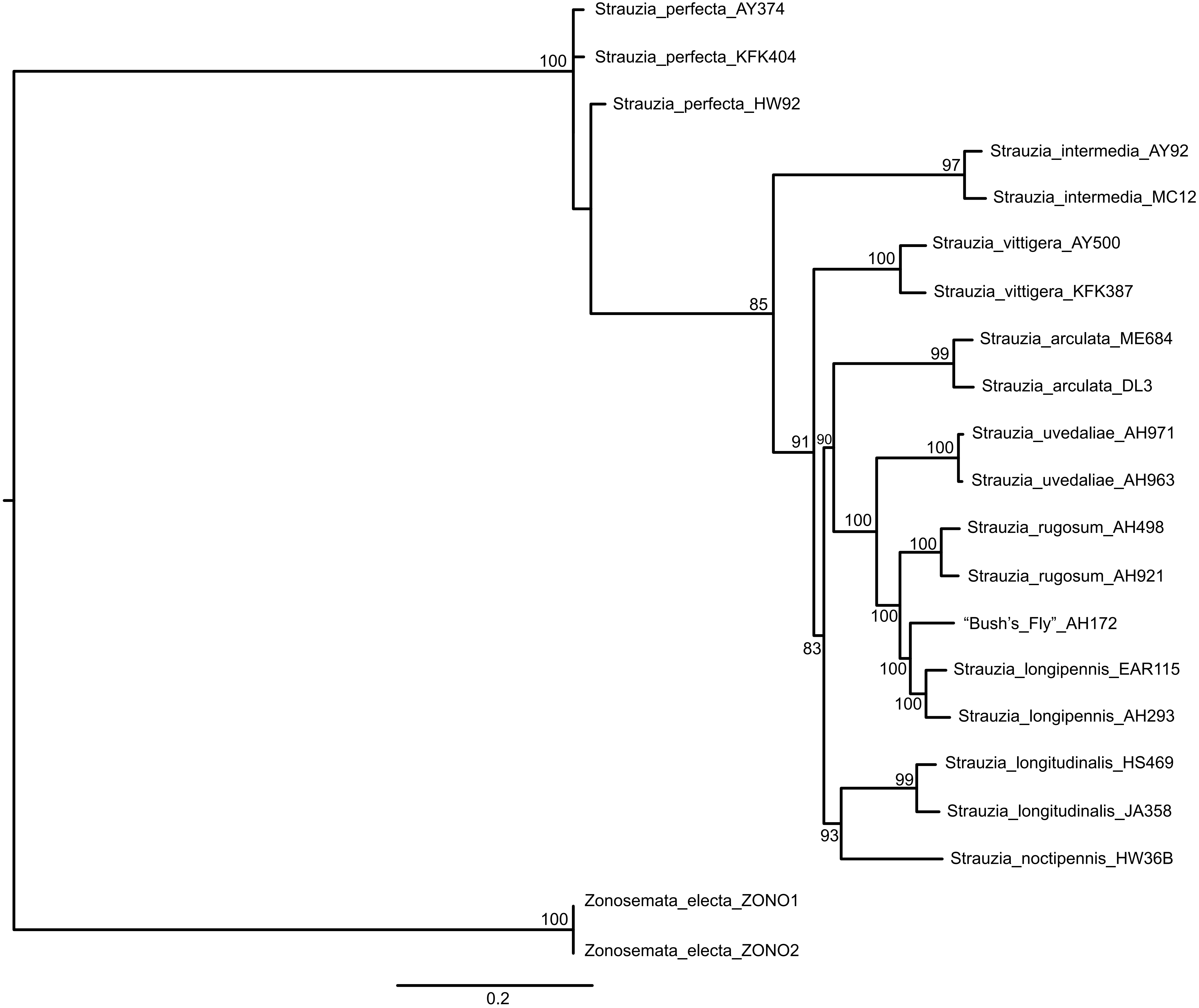

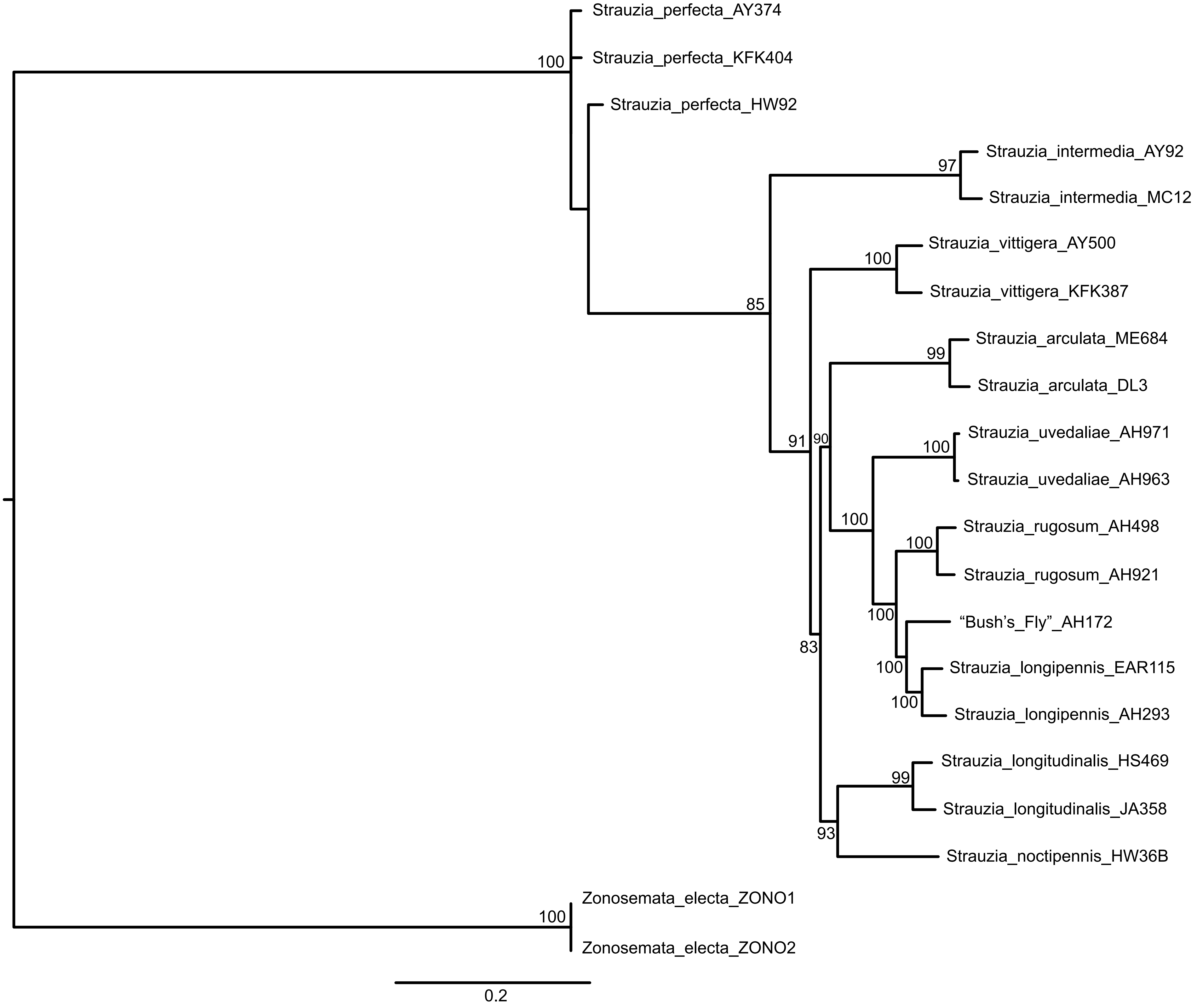

### Supplemental Figure 4

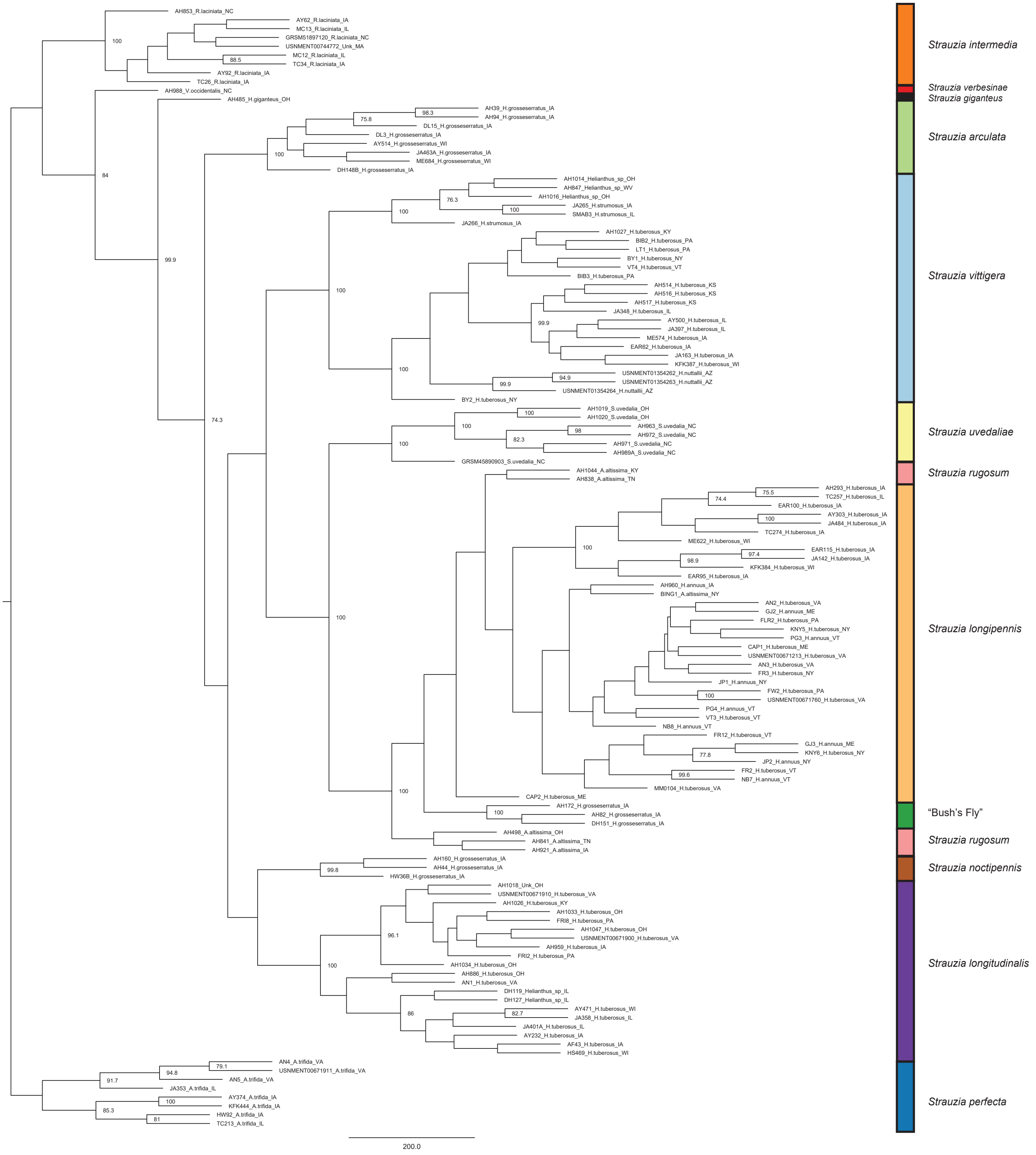

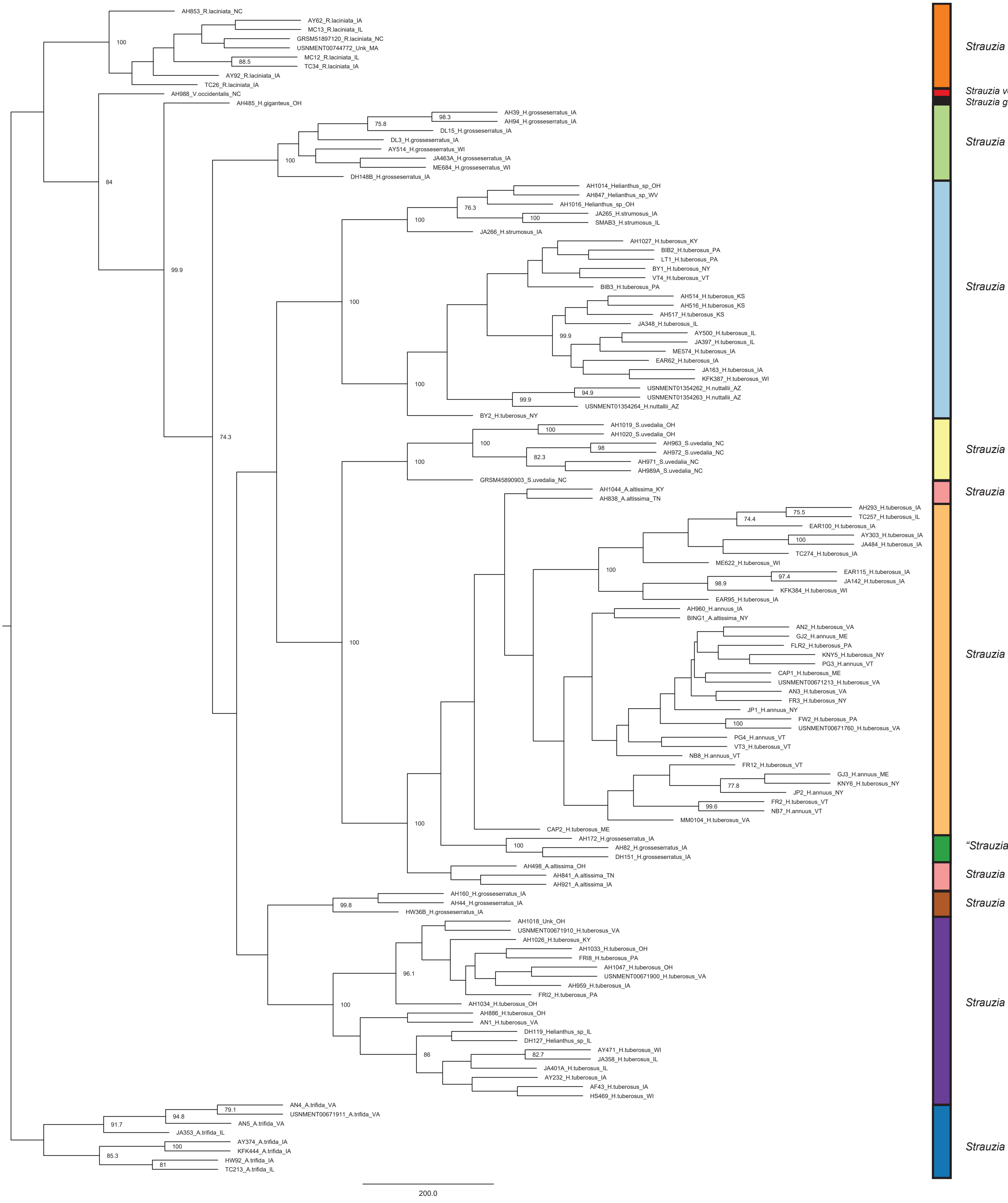
