## Supplemental Table 2 Page 1 for "Host shifting and host sharing in a genus of specialist flies diversifying alongside their sunflower hosts"

**Sequencing Project 1:**

| Sample ID | i7 adapter | | Forward Barcode | Reverse Barcode |
|---|---|---|---|---|
| AF43 | Tru7 01-06 | GCTGTAAG | TCCGAATG | TTCGGTACT |
| AH160 | Tru7 01-08 | TTCGGCTA | TCTGCAACTG | TTGTGCACGAT |
| AH172 | Tru7 01-08 | TTCGGCTA | TAGCGTTGG | TTGTGCACGAT |
| AH293 | Tru7 01-06 | GCTGTAAG | TGATACCG | TTGTGCACGAT |
| AH39 | Tru7 01-06 | GCTGTAAG | TGGTCTACGTG | TGCATCAT |
| AH44 | Tru7 01-06 | GCTGTAAG | TTTAGGCAG | TTCGGTACT |
| AH514 | Tru7 01-06 | GCTGTAAG | TCCGAATG | TGCATCAT |
| AH516 | Tru7 01-06 | GCTGTAAG | TTCATGGTCAG | TTGTGCACGAT |
| AH517 | Tru7 01-06 | GCTGTAAG | TCTGCAACTG | TATGCTGTT |
| AH82 | Tru7 01-06 | GCTGTAAG | TCTGCAACTG | TCTAACGT |
| AH853 | Tru7 01-08 | TTCGGCTA | TCCGAATG | TCACATGTCT |
| AH886 | Tru7 01-06 | GCTGTAAG | TCTGCAACTG | TCATGACCTT |
| AH921 | Tru7 01-06 | GCTGTAAG | TGATACCG | TCATGACCTT |
| AH94 | Tru7 01-06 | GCTGTAAG | TGGTCTACGTG | TATGCTGTT |
| AN1 | Tru7 01-08 | TTCGGCTA | TTTAGGCAG | TTGTGCACGAT |
| AY232 | Tru7 01-01 | AGTGACCT | TGATACCG | TACGCATT |
| AY303 | Tru7 01-08 | TTCGGCTA | TTCATGGTCAG | TCACATGTCT |
| AY374 | Tru7 01-08 | TTCGGCTA | TAGCGTTGG | TCACATGTCT |
| AY471 | Tru7 01-06 | GCTGTAAG | TCCGAATG | TCATGACCTT |
| AY500 | Tru7 01-06 | GCTGTAAG | TCTGCAACTG | TGCATCAT |
| AY514 | Tru7 01-06 | GCTGTAAG | TCCGAATG | TATGCTGTT |
| AY62 | Tru7 01-08 | TTCGGCTA | TTCATGGTCAG | TTGTGCACGAT |
| AY92 | Tru7 01-06 | GCTGTAAG | TAACTCGTCG | TTCGGTACT |
| DH119 | Tru7 01-06 | GCTGTAAG | TAACTCGTCG | TTGTGCACGAT |
| DH127 | Tru7 01-08 | TTCGGCTA | TGGTCTACGTG | TCACATGTCT |
| DH148B | Tru7 01-06 | GCTGTAAG | TCCGAATG | TCTAACGT |
| DH151 | Tru7 01-06 | GCTGTAAG | TTTAGGCAG | TGCATCAT |
| DL15 | Tru7 01-01 | AGTGACCT | TGGTCTACGTG | TAGCTACACTT |
| DL3 | Tru7 01-06 | GCTGTAAG | TAGCGTTGG | TGCATCAT |
| EAR100 | Tru7 01-01 | AGTGACCT | TTCATGGTCAG | TTGTGCACGAT |
| EAR115 | Tru7 01-06 | GCTGTAAG | TTTAGGCAG | TATGCTGTT |
| EAR62 | Tru7 01-06 | GCTGTAAG | TAACTCGTCG | TCATGACCTT |
| EAR95 | Tru7 01-06 | GCTGTAAG | TGATACCG | TGCATCAT |
| HS469 | Tru7 01-06 | GCTGTAAG | TTCATGGTCAG | TTCGGTACT |
| HW36B | Tru7 01-01 | AGTGACCT | TCCGAATG | TCTAACGT |
| HW92 | Tru7 01-06 | GCTGTAAG | TCTGCAACTG | TTCGGTACT |
| JA142 | Tru7 01-06 | GCTGTAAG | TAACTCGTCG | TATGCTGTT |
| JA163 | Tru7 01-06 | GCTGTAAG | TTCATGGTCAG | TCTAACGT |
| JA266 | Tru7 01-06 | GCTGTAAG | TTCATGGTCAG | TGCATCAT |
| JA348 | Tru7 01-06 | GCTGTAAG | TGATACCG | TATGCTGTT |
| JA353 | Tru7 01-06 | GCTGTAAG | TTTAGGCAG | TCATGACCTT |
| JA358 | Tru7 01-06 | GCTGTAAG | TAGCGTTGG | TTGTGCACGAT |
| JA397 | Tru7 01-06 | GCTGTAAG | TAGCGTTGG | TTCGGTACT |
| JA401A | Tru7 01-06 | GCTGTAAG | TGGTCTACGTG | TCTAACGT |
| JA463A | Tru7 01-06 | GCTGTAAG | TCCGAATG | TTGTGCACGAT |
| JA484 | Tru7 01-01 | AGTGACCT | TCTGCAACTG | TCACATGTCT |
| KFK384 | Tru7 01-06 | GCTGTAAG | TAACTCGTCG | TGCATCAT |
| KFK387 | Tru7 01-06 | GCTGTAAG | TTTAGGCAG | TCTAACGT |
| KFK444 | Tru7 01-08 | TTCGGCTA | TCTGCAACTG | TCACATGTCT |
| MC12 | Tru7 01-06 | GCTGTAAG | TAGCGTTGG | TCTAACGT |
| MC13 | Tru7 01-08 | TTCGGCTA | TGATACCG | TCACATGTCT |
| ME574 | Tru7 01-08 | TTCGGCTA | TCCGAATG | TTGTGCACGAT |
| ME622 | Tru7 01-06 | GCTGTAAG | TGATACCG | TCTAACGT |
| ME684 | Tru7 01-06 | GCTGTAAG | TTTAGGCAG | TTGTGCACGAT |
| MM0104 | Tru7 01-08 | TTCGGCTA | TAACTCGTCG | TTGTGCACGAT |
| TC213 | Tru7 01-06 | GCTGTAAG | TGGTCTACGTG | TTCGGTACT |
| tc257 | Tru7 01-06 | GCTGTAAG | TGGTCTACGTG | TTGTGCACGAT |
| TC26 | Tru7 01-06 | GCTGTAAG | TAACTCGTCG | TCTAACGT |
| TC274 | Tru7 01-06 | GCTGTAAG | TGATACCG | TTCGGTACT |
| TC34 | Tru7 01-01 | AGTGACCT | TAGCGTTGG | TGTATGCAT |
| ZONO1 | Tru7 01-08 | TTCGGCTA | TGGTCTACGTG | TTGTGCACGAT |
| ZONO2 | Tru7 01-08 | TTCGGCTA | TGATACCG | TTGTGCACGAT |
