## Supplemental Table 2 Page 2 for "Host shifting and host sharing in a genus of specialist flies diversifying alongside their sunflower hosts"

**Sequencing Project 2:**

| **Sample ID** | **i7 adapter** | | **Forward Barcode** | **Reverse Barcode** |
|---|---|---|---|---|
| AH1014 | Tru7 01-06 | GCTGTAAG | TTCATGGTCAG | TATGCTGTT |
| AH1016 | iTru7 01-08 | TTCGGCTA | TGGTCTACGTG | TAGCTACACTT |
| AH1018 | iTru7 01-08 | TTCGGCTA | TGGTCTACGTG | TACGCATT |
| AH1019 | Tru7 01-06 | GCTGTAAG | TAGCGTTGG | TATGCTGTT |
| AH1020 | iTru7 01-08 | TTCGGCTA | TCCGAATG | TACGCATT |
| AH1026 | Tru7 01-06 | GCTGTAAG | TGGTCTACGTG | TCATGACCTT |
| AH1027 | iTru7 01-08 | TTCGGCTA | TTCATGGTCAG | TAGCTACACTT |
| AH1033 | Tru7 01-06 | GCTGTAAG | TCCGAATG | TTGCAGTGAGT |
| AH1034 | iTru7 01-08 | TTCGGCTA | TTTAGGCAG | TACGCATT |
| AH1044 | Tru7 01-06 | GCTGTAAG | TAGCGTTGG | TCATGACCTT |
| AH1047 | Tru7 01-06 | GCTGTAAG | TAACTCGTCG | TCATGACCTT |
| AH485 | Tru7 01-06 | GCTGTAAG | TCTGCAACTG | TATGCTGTT |
| AH498 | iTru7 01-08 | TTCGGCTA | TCTGCAACTG | TAGCTACACTT |
| AH838 | iTru7 01-08 | TTCGGCTA | TAGCGTTGG | TAGCTACACTT |
| AH841 | Tru7 01-06 | GCTGTAAG | TGATACCG | TATGCTGTT |
| AH847 | iTru7 01-08 | TTCGGCTA | TAGCGTTGG | TACGCATT |
| AH959 | itru7 01-08 | TTCGGCTA | TTTAGGCAG | TTCGGTACT |
| AH960 | Tru7 01-06 | GCTGTAAG | TCTGCAACTG | TGTATGCAT |
| AH963 | Tru7 01-06 | GCTGTAAG | TGATACCG | TCATGACCTT |
| AH971 | Tru7 01-06 | GCTGTAAG | TGGTCTACGTG | TCACATGTCT |
| AH972 | itru7 01-08 | TTCGGCTA | TGATACCG | TCTAACGT |
| AH988 | Tru7 01-06 | GCTGTAAG | TTTAGGCAG | TTGCAGTGAGT |
| AH989A | Tru7 01-06 | GCTGTAAG | TCTGCAACTG | TCATGACCTT |
| AN2 | Tru7 01-06 | GCTGTAAG | TCCGAATG | TATGCTGTT |
| AN3 | iTru7 01-08 | TTCGGCTA | TTTAGGCAG | TAGCTACACTT |
| AN4 | Tru7 01-06 | GCTGTAAG | TTTAGGCAG | TATGCTGTT |
| AN5 | iTru7 01-08 | TTCGGCTA | TAACTCGTCG | TAGCTACACTT |
| BIB2 | Tru7 01-06 | GCTGTAAG | TGGTCTACGTG | TATGCTGTT |
| BIB3 | iTru7 01-08 | TTCGGCTA | TCCGAATG | TAGCTACACTT |
| BING1 | Tru7 01-06 | GCTGTAAG | TCCGAATG | TCATGACCTT |
| BY1 | Tru7 01-06 | GCTGTAAG | TTTAGGCAG | TGTATGCAT |
| BY2 | itru7 01-08 | TTCGGCTA | TCTGCAACTG | TCTAACGT |
| CAP1 | Tru7 01-06 | GCTGTAAG | TAACTCGTCG | TGTATGCAT |
| CAP2 | Tru7 01-06 | GCTGTAAG | TCTGCAACTG | TGCATCAT |
| FLR2 | Tru7 01-06 | GCTGTAAG | TTTAGGCAG | TCATGACCTT |
| FR12 | itru7 01-08 | TTCGGCTA | TAACTCGTCG | TCTAACGT |
| FR2 | Tru7 01-06 | GCTGTAAG | TAGCGTTGG | TGTATGCAT |
| FR3 | Tru7 01-06 | GCTGTAAG | TGGTCTACGTG | TGCATCAT |
| FRI2 | Tru7 01-06 | GCTGTAAG | TAACTCGTCG | TATGCTGTT |
| FRI8 | iTru7 01-08 | TTCGGCTA | TGATACCG | TAGCTACACTT |
| FW2 | Tru7 01-06 | GCTGTAAG | TTTAGGCAG | TCACATGTCT |
| GJ2 | Tru7 01-06 | GCTGTAAG | TGATACCG | TGTATGCAT |
| GJ3 | itru7 01-08 | TTCGGCTA | TTCATGGTCAG | TCTAACGT |
| GRSM45890903 | Tru7 01-06 | GCTGTAAG | TGGTCTACGTG | TGTATGCAT |
| GRSM51897120 | Tru7 01-06 | GCTGTAAG | TTTAGGCAG | TTGTGCACGAT |
| JA265 | itru7 01-08 | TTCGGCTA | TTTAGGCAG | TGATCGTTGT |
| JP1 | Tru7 01-06 | GCTGTAAG | TTCATGGTCAG | TCACATGTCT |
| JP2 | itru7 01-08 | TTCGGCTA | TGGTCTACGTG | TCTAACGT |
| KNY5 | Tru7 01-06 | GCTGTAAG | TCCGAATG | TTGTGCACGAT |
| KNY6 | itru7 01-08 | TTCGGCTA | TTTAGGCAG | TCTAACGT |
| LT1 | Tru7 01-06 | GCTGTAAG | TCCGAATG | TCACATGTCT |
| NB7 | Tru7 01-06 | GCTGTAAG | TCCGAATG | TGTATGCAT |
| NB8 | Tru7 01-06 | GCTGTAAG | TCCGAATG | TGCATCAT |
| PG3 | Tru7 01-06 | GCTGTAAG | TAACTCGTCG | TCACATGTCT |
| PG4 | itru7 01-08 | TTCGGCTA | TCCGAATG | TCTAACGT |
| SMAB3 | itru7 01-08 | TTCGGCTA | TAGCGTTGG | TTCGGTACT |
| USNMENT00671213 | Tru7 01-06 | GCTGTAAG | TGATACCG | TCACATGTCT |
| USNMENT00671760 | itru7 01-08 | TTCGGCTA | TTCATGGTCAG | TTCGGTACT |
| USNMENT00671900 | Tru7 01-06 | GCTGTAAG | TTTAGGCAG | TGCATCAT |
| USNMENT00671910 | Tru7 01-06 | GCTGTAAG | TTCATGGTCAG | TGTATGCAT |
| USNMENT00671911 | itru7 01-08 | TTCGGCTA | TAGCGTTGG | TCTAACGT |
| USNMENT00744772 | Tru7 01-06 | GCTGTAAG | TAACTCGTCG | TTGTGCACGAT |
| USNMENT01354262 | Tru7 01-06 | GCTGTAAG | TCTGCAACTG | TCACATGTCT |
| USNMENT01354263 | Tru7 01-06 | GCTGTAAG | TAACTCGTCG | TGCATCAT |
| USNMENT01354264 | itru7 01-08 | TTCGGCTA | TAACTCGTCG | TTCGGTACT |
| VT3 | itru7 01-08 | TTCGGCTA | TCTGCAACTG | TTCGGTACT |
| VT4 | Tru7 01-06 | GCTGTAAG | TAGCGTTGG | TCACATGTCT |
